# Molecular basis of neurodevelopmental disorder-causing mutation in nonsense-mediated mRNA decay factor UPF3B

**DOI:** 10.1101/2022.04.03.486873

**Authors:** Joshua C. Bufton, Kyle T. Powers, Jenn-Yeu A. Szeto, Christiane Schaffitzel

## Abstract

UPF3B is a key nonsense-mediated mRNA decay (NMD) factor required for surveillance of mRNA and regulation of eukaryotic gene expression. Mutations in UPF3B cause intellectual disability. The underlying molecular mechanisms remain unexplored as the mutations lie in an uncharacterized region of UPF3B. Here, we show that UPF3B shares structural and functional homology to the Drosophila Behavior/Human Splicing protein family comprising an RNA-recognition motif-like domain (RRM-L), a NONA/paraspeckle-like domain (NOPS-L), and extended α-helical domains essential for ribosome- and RNA-binding and RNA-induced oligomerization. A co-crystal structure of UPF3B with the third middle domain of eukaryotic initiation factor 4G (MIF4GIII) of UPF2 reveals an unexpectedly intimate binding interface. UPF3B’s disease-causing mutation Y160D located in the NOPS-L domain reduces the UPF2 binding affinity ~40-fold compared to wildtype UPF3B. UPF3B’s paralogue UPF3A, an NMD antagonist which is upregulated in patients with the UPF3B-Y160D mutation, binds UPF2 with ~10-fold higher affinity than UPF3B, leading to impaired NMD activity and upregulation of mRNAs involved in neurodevelopment.

## INTRODUCTION

The nonsense-mediated mRNA decay (NMD) pathway targets mRNAs harboring premature termination codons (PTCs) thus preventing translation of potentially toxic C-terminally truncated proteins (1–5). NMD is clinically highly relevant because nonsense mutations account for ~20% of known disease-associated single base-pair substitutions (6). Simultaneously, NMD also has a conserved and fundamental role in non-aberrant eukaryotic gene expression, for example in regulating neurodevelopment in mammals (7–9). The NMD machinery comprises the conserved UP-Frameshift proteins UPF1, UPF2 and UPF3B (10–12). UPF3B is an auxiliary component of exon-junction complexes (EJC) that are deposited at exon-exon boundaries during mRNA splicing and act as enhancers of mammalian NMD (13,14). The interaction between the UPF proteins is critical for NMD progression. The UPF2-UPF3B complex binds to UPF1 leading to structural rearrangements and stimulation of ATPase and helicase activity of UPF1 (15,16). In addition, UPF2-UPF3B is suggested to activate SMG1 kinase-mediated UPF1 phosphorylation initiating mRNA decay by recruitment of the nucleases (17,18). More recently mRNA and ribosome binding of UPF3B has been reported (19,20). UPF3B was shown to slow down translation termination and support ribosome dissociation *in vitro* (19). Cross-linking studies showed UPF3B binding to mRNA upstream of exon-exon junctions (20) suggesting a role of UPF3B in the positioning of EJCs and the NMD machinery. The precise mode of action of UPF3B in NMD remains enigmatic due to the absence of structural and functional data. UPF3B’s N-terminus comprises a conserved RNA recognition motif-like domain (RRM-L) lacking residues essential for RNA interaction (21–23) (Figure 1A). Instead, the RRM-L binds the third middle domain of eukaryotic initiation factor 4G (MIF4GIII) of UPF2 (12,21,24). The RRM-L domain is followed by an uncharacterized middle-domain and a C-terminal EJC-binding motif (Figure 1A).

**Figure 1.**
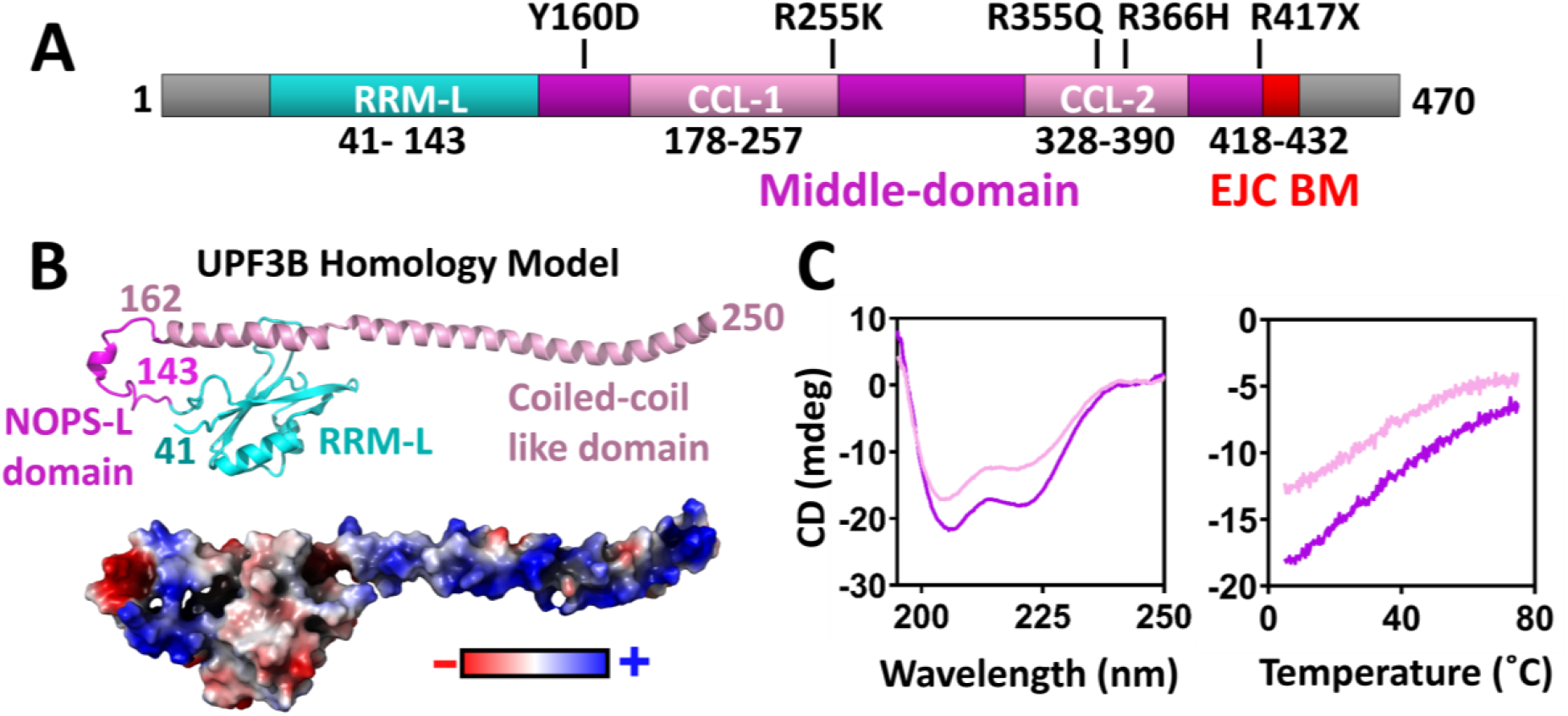
UPF3B domain architecture and homology with DBHS proteins. **(A)** UPF3B isoform2 domain organization highlighting known disease-causing single base pair mutations. **(B)** Above: UPF3B homology model created with MODELLER using the structure of DBHS protein SFPQ (PDB ID: 4WIK) as a template (37) (above). UPF3B’s RRM-L is colored in cyan, predicted NOPS-like domain (NOPS-L) in magenta and coiled-coil-like domain in light pink. Below: Surface representation of the UPF3B homology model indicating electrostatic potential; positive (blue), neutral (white) and negative (red). The map was produced using the Adaptive Poisson-Boltzmann Solver within PyMol. **(C)** Left: CD spectroscopy trace of a 195-250 nm wavelength scan for UPF3B-146-256 (light pink line) and UPF3B-146-417 (magenta line) truncations revealing a double dip trace indicative of α-helical structure. Right: temperature melt of the same constructs from 5-75°C taking CD measurements at 222 nm indicating a non-sigmoidal curve.

UPF3B missense mutations were identified in patients with schizophrenia and X-linked intellectual disability (XLID) (25–29) (Figure 1A). Expression of these UPF3B mutants in neural stem cells impairs neuronal differentiation and reduces neurite branching (25). Specifically, mutations Y160D and R366H, located in the uncharacterized middle domain of UPF3B result in subtly increased expression of NMD targets ARHGAP24 and ATF4 which are involved in down-regulation of neuronal branching and plasticity (25,30,31). The disease-causing UPF3B-Y160D mutation additionally leads to substantial upregulation of UPF3A, a paralogue of UPF3B which can protect itself from degradation by interaction with UPF2 (28,32). UPF3A is a weaker NMD activator and can partially compensate for UPF3B loss, but is also reported to antagonize NMD (14,28,32–35). Tyrosine residue Y160 of UPF3B is highly conserved in vertebrates, *Drosophila melanogaster*, *Caenorhabditis elegans* and also in vertebrate UPF3A (25).

Here, we report the co-crystal structure of UPF3B-UPF2, define the residues involved in high-affinity interactions with UPF2 and map UPF3B’s nucleic acid binding domains. We show that the middle-domain of UPF3B adopts a NONA/paraspeckle-like (NOPS-L) fold followed by extended α-helical domains which are structurally homologous to paraspeckle proteins. In agreement with this homology, we observe RNA- and DNA-induced oligomerization of UPF3B. UPF3B’s binding of RNA and DNA is promiscuous, with preference for double-stranded RNA which is prominent in ribosomes. We map nucleic acid-binding and oligomerization activities onto the RRM-L and the N-terminal part of the middle domain, encompassing the NOPS-L and first α-helical domain. A co-crystal structure of UPF3B comprising these domains with UPF2-MIF4GIII reveals an unexpectedly intricate binding interface. UPF2-MIF4GIII is wedged between UPF3B’s RRM-L and NOPS-L, and the newly identified interactions between the middle-domain and UPF2 lead to a >200-fold affinity increase compared to UPF2-MIF4GIII and the RRM-L domain only. UPF3B residue Y160 and adjacent conserved hydrophobic residues of the NOPS-L domain bind into a hydrophobic cleft of UPF2. Accordingly, the UPF3B-Y160D mutation implicated in XLID reduces UPF3B’s affinity for UPF2-MIF4GIII ~40-fold. UPF3A-isoform1, which is upregulated in patients with the UPF3B-Y160D mutation, binds UPF2-MIF4GIII with even higher, picomolar affinity indicating an additional level of regulation of UPF2-UPF3 interaction rather than a direct competition between UPF3A and UPF3B for UPF2 binding in cells.

## MATERIALS AND METHODS

### Secondary structure prediction, homology modelling, and sequence conservation analysis

Secondary structure predictions were generated using the primary amino acid sequence of UPF3B isoform2 and submitted to Quick2D as well as PCOILS, MARCOIL, and Deepcoil servers within the MPI Bioinformatics Toolkit (36). Highest scoring hits were considered those with a >95% probability of homology despite having 10-20% sequence identity with UPF3B including structures derived from splicing regulation: *Schizosaccharomyces pombe* Mei2, *Mus musculus* TIA-1, *Saccharomyces cerevisiae* PRP24, *Homo sapiens* RBM5, and ribosomal biogenesis *S. cerevisiae* NOP15 (PDB IDs: 6YYL, 2DGO, 2L9W, 2LKZ and 3JCT_o respectively). Homology modelling of UPF3B was conducted using HHPRED and MODELLER (37) servers using SFPQ PDB ID 4WIK to build the putative domains of UPF3B’s middle region. The WebLogo server was used to generate sequence conservation across the putative structured middle domain of UPF3B (38).

### Construct design and cloning

pFastBac-HTB (Invitrogen) baculoviral expression vectors for UPF3B-FL (residues 1-470), UPF3B-M (146-417) and UPF3B-SM (146-256) were reported previously (19). UPF3B constructs UPF3B-41-143, UPF3B-41-189, UPF3B-41-214, UPF3B-41-262, and UPF2-MIF4GIII (761-1054), UPF2L (120-1227) were produced by PCR amplification (primers listed in Supplementary Table S1) using the Phusion High-Fidelity PCR kit (NEB #E0553S) prior to restriction digest and ligation into pPROEX-HTB (Invitrogen) vectors. UPF3B-41-189 derived point mutants including Y160A, Y160D, Y167A, Y167D, Y160A+Y167A, Y160D+Y167D and Avi-tagged Avi-UPF2-MIF4GIII as well as Avi-UPF3B-WT constructs were generated using a Q5 Site-Directed Mutagenesis Kit (NEB #E0554S) with mutagenic primers (Eurofins Genomics) (Supplementary Table S2). Fragments for UPF3A-58-206-Iso1 and UPF3A-58-173-Iso2 were codon-optimized for *Escherichia coli* expression and gene-synthesized (Twist Bioscience) prior to restriction digest and ligation into pPROEX-HTB (Invitrogen) vector.

### Protein expression and purification

UPF3B-FL, Avi-UPF3B-WT, UPF3B-M, and UPF3B-SM were expressed using the MultiBac insect cell expression system (39). All other UPF3B, UPF3A, and UPF2 (MIF4GIII + UPF2L) constructs were transformed into *E. coli* BL21 Rosetta2(DE3) (Novagen #71400) and grown in LB medium prior to induction with 0.3 mM isopropyl 1-thio-β-D galactopyranoside (IPTG) and expressed for 16 hours at 18 °C with agitation. Cells were harvested by centrifugation at 1000 x g and 5000 x g for insect and bacterial cells respectively, before flash freezing pellets for storage at −80 °C.

For purification, cell pellets were thawed prior to resuspension in binding buffer (25 mM HEPES pH 7.4, 10 mM imidazole, 300 mM NaCl, 5% [v/v] glycerol) supplemented with 1x cOmplete EDTA free protease inhibitor cocktail tablet (Roche #11873580001). Cells were lysed via sonication, and the lysate clarified at 45,000 x g for 45 min at 4° C. The supernatant was applied onto a 5 mL HiFliQ Ni-NTA column (Generon #HiFliQ5-NiNTA-5). After washing the column with binding buffer, bound 6xHis-proteins were eluted via a linear gradient of 350 mM imidazole (in binding buffer). Eluted protein-containing fractions were incubated with in-house purified Tobacco Etch Virus (TEV) protease for removal of the 6xHis-tag. All proteins were further purified by reverse IMAC to remove un-cleaved material prior to tandem ion exchange purification using a 5mL HiTrap Q XL (Cytiva #17515901) column followed by a second cation exchange 5 mL HiTrap SP HP column (Cytiva #17115201). After loading the sample, the Q XL column was removed and elution carried out on SP HP column using a linear gradient from 150 mM to 1 M NaCl in 25 mM HEPES pH 7.4, 2 mM β-mercaptoethanol. For UPF2-MIF4GIII and UPF2L constructs, ion exchange chromatography was carried out using the SP HP column only. Proteins were concentrated via Amicon centrifugal ultrafiltration units with a MWCO of 3 kDa. Protein concentrations were determined by measuring UV absorbance at 280 nm with calculated molecular weights and extinction coefficients using a NanoDrop One (ThermoFisher).

### Electrophoretic mobility shift assays (EMSA)

For UPF3B-nucleic acid interaction analysis, HEX-labelled oligos (Supplementary Table S1, Eurofins Genomics) were diluted to 2 μM (2x 1 μM final) in EMSA buffer (25 mM HEPES pH 7.45, 2 mM MgCl2, 150 mM NaCl, 0.5 mM TCEP). 1:1 serial dilutions of UPF3B-WT were then added to the nucleotide solutions and incubated on ice for 30 mins before supplementing with 7.5% (final) glycerol and running on a Novex 6% Tris-Glycine WedgeWell gel (Invitrogen #XP0006A) equilibrated in Novex Tris-Glycine Native Running Buffer at 4 °C. Gels were visualized with a Typhoon FLA 9500 (GE Healthcare) imager with an excitation wavelength of 532 nm using a BPG1 emission filter. For UPF3B-UPF2-ssRNA interactions, ssRNA was diluted to 2 μM in EMSA buffer along with either 2 μM UPF3B-WT or UPF2L and increasing ratios of either UPF2L or UPF3B. Incubations were performed on ice for 1 hour before loading on a 4 to 20% Novex WedgeWell Tris-Glycine native gel prior to staining with Coomassie blue stain.

### Circular dichroism (CD) spectroscopy assays

All CD measurements were performed in a 0.1 cm quartz cuvette using a J-1500 spectropolarimeter (Jasco) fitted with a Peltier temperature control unit. An initial CD wavelength scan measurement at 25 °C was carried out with 10 μM of UPF3B-146-256 and UPF3B-146-417 dialyzed in CD buffer (100 mM KCl, 10 mM potassium phosphate pH 7.4). CD spectra were acquired across a wavelength range of 195-260 nm collecting data at 0.1 nm intervals. For the temperature wavelength scan, CD measurements at 222 nm were recorded at 0.2 °C intervals from 5 to 75 °C with heating at 5 °C/min and a 10 second equilibration time at each temperature point. A buffer-only derived baseline was subtracted from all datasets.

### Fluorescence anisotropy (FA) assays

HEX-labelled DNA (5’ HEX-CCCTGAGCTGACGCAGCACCTGGG 3’) and RNA probes (5’ CCCUGAGCUGACGCAGCACCUGGG 3’) (Eurofins Genomics) were diluted to 5 nM in assay buffer (150 mM NaCl, 25 mM HEPES pH 7.45, 1 mM TCEP). An assay volume of 150 μl was dispensed into a Hellma 10 x 2 mm Suprasil quartz cuvette (Merck #Z802778). For double-stranded substrates, complementary oligonucleotides were mixed at a 1:1 molar ratio and heated to 65 °C prior to slowly cooling to room temperature overnight in an insulated chamber. Proteins were then titrated into the cuvette at increasing concentrations and fluorescence anisotropy measurements were recorded at 20 °C with a Jobin Yvon Fluorolog (Horiba Scientific) with excitation and emission wavelengths of 530 nm and 550 nm respectively. Measurements were taken with an integration time of 0.5 seconds and averaged across 4 accumulations. Dilutions and titrations were carried out in triplicate and averages with standard deviations were plotted before fitting the data (GraphPad Prism) with a single-component binding equation 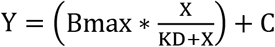 to determine the dissociation binding constant (K_D_) where B_max_ is the maximum anisotropy change (saturated anisotropy value - starting value), X is the concentration of protein, and C is the anisotropy value at which the X axis =0.

### Surface plasmon resonance (SRP) assays

SPR experiments were performed on a Biacore T200 using a SA-Series S sensor chip (Cytiva #BR100531). Avi-UPF2-MIF4GIII and Avi-UPF3B-WT were biotinylated via incubation with BirA as previously described (40) prior to size exclusion chromatography to remove free biotin. Immobilization of biotinylated ligands was carried out on a single flow cell leaving a 2^nd^ flow cell as a background control for signal subtraction and all analysis was performed at 15 °C. For UPF2-UPF3 interaction analysis, 350 RU of biotinylated-Avi-UPF2-MIF4GIII was immobilized. UPF3B and UPF3A construct serial dilutions in Biacore running buffer (300 mM NaCl, 25 mM HEPES pH 7.4, 0.25 mM TCEP, 0.05% Tween-20) were injected at 30 μL/min with an association phase of 180 s and a dissociation phase of 240 s. After every dissociation, the surface was regenerated with a 120 s injection of regeneration solution (1 M MgCl_2_, 25 mM HEPES pH 7.4, 0.05% Tween-20, 5% glycerol). Resulting sensorgrams were analyzed with the Biacore evaluation software (version 1.0) yielding on- and off-rates obtained through a global fit of both the association and dissociation phases of at least four different concentrations of each analyte using the 1:1 binding model. For testing ribosome subunit binding, 150 RU of biotinylated Avi-UPF3B (Full length WT) was immobilized. A dilution series of 40S/ 60S subunits was made in ribosomal suspension buffer (200 mM KCl, 2.5 mM Mg(OAc)_2_, 20 mM HEPES pH 7.6, 0.5 mM TCEP, 10 mM NH4Cl, 0.05% Tween-20) and injected at 20 μL/min for 6 min followed by a 12 min dissociation phase. The surface was regenerated by two 90 s injections of regeneration solution (1M KCl, 20 mM HEPES pH 7.4, 0.05% Tween-20, 20 mM EDTA) prior to the next injection. Data was analyzed by steady-state fit in the Biacore evaluation software by measuring the RU values of each concentration 20 s before the end of the association phase. Equilibrium dissociation constants (K_D_) were determined by plotting the RU values as a function of analyte concentration and fitted with a single-component binding equation in GraphPad Prism 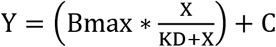. Reported K_D_ values are an average of three or more experimental repeats with ranges reported by standard deviation.

### Protein crystallization

We performed crystallization experiments of various UPF3B truncations alone, with RNA oligomers, and with UPF2-MIF4GIII. The latter was mixed at a 1:1 molar ratio with UPF3B truncation constructs encompassing RRM-L and parts of the middle domain (UPF3B-41-189, UPF3B-41-214, and UPF3B-41-262) prior to purification via size-exclusion chromatography (SEC) using a Superdex 200 column (Cytiva #17517501) equilibrated with GF buffer (25 mM HEPES pH 7.4, 300 mM NaCl, 2 mM TCEP). Crystallization experiments were carried out using the standard sitting drop vapor diffusion method on LOW PROFILE Swissci Polystyrene Triple Drop Plates (Molecular Dimensions # MD11-003LP-100) containing 45 μL of reservoir solution. Using a mosquito crystal (SPT Labtech), 125, 150 and 175 nL of UPF3B-41.189+UPF2-MIF4GIII at an equimolar complex concentration of 15 mg/mL in GF buffer was dispensed along with 175, 150 and 125 nL of reservoir solution in drops A, B and C respectively resulting in 300 nL total drop volume. Plates were incubated at both 4 °C and 20 °C.

### Data collection and structure determination

We had several crystallization hits for the UPF3B-41-189 construct with UPF2-MIF4GIII in the initial screens. The best diffracting crystals appeared after 7 days at 20 °C in 0.1 M calcium acetate, 0.1 M MES pH 6.0, 15% PEG 400 and continued to grow over the course of two weeks. Drops were supplemented with 20% Ethylene Glycol (final concentration) upon harvesting as a cryoprotectant prior to flash freezing in liquid nitrogen. Diffraction data were collected on the IO4-1 beamline at the Diamond Light Source (Harwell Science and Innovation Campus) and images processed using XDS (Version: 01/2020) and scaled with AIMLESS (41,42). The crystals diffracted to a resolution of 2.6 Å in space group P4_1_22 with four complexes in the asymmetric unit. The structure was phased by molecular replacement in PHASER using the previously solved UPF3B-RRM+UPF2-MIF4GIII (PDB ID:1UW4) as an input model (21). The crystallographic asymmetric unit was found to be different to the previously determined structure containing four UPF2-MIF4GIII-UPF3B complexes rather than two, forming different crystal packing contacts and in the space group of P4_1_22 rather than P2_1_2_1_2. The four copies of UPF3B-UPF2-MIF4GIII heterodimers in the asymmetric unit were found to be very similar having an average RMSD of 0.3 Å between complexes (Table 1). Manual model building was performed in Coot (version 0.8.9) and automated refinement carried out in PHENIX.REFINE (PHENIX version 1.17.1-3660) (43,44). Models were validated and statistics obtained using MolProbity (45). Figures were prepared using PyMol (version 2.3.4).

**Table 1.** Refinement statistics of UPF3B-41-189 + UPF2-MIF4GIII co-crystal structure.

| Data collection |  |  |  |
| --- | --- | --- | --- |
| Wavelength (Å) | 0.9795 |  |  |
| Space group | $P4_122$ | | |
| Cell Dimensions |  |  |  |
| $a, b, c$ (Å) | 130.59 | 130.59 | 267.33 |
| $\alpha, \beta, \gamma$ (°) | 90.00 | 90.00 | 90.00 |
| Collection Statistics |  |  |  |
| Resolution range (Å) | 1.309 - 30 |  |  |
| Completeness (%) | 99.94 |  |  |
| $R_{\text{pim}}$ (Highest Shell) | 0.061 (0.809) | | |
| $I/\sigma I$ (Highest Shell) | 10.2 (1.0) | | |
| Total reflections | 1173009 |  |  |
| Unique reflections | 80565 |  |  |
| Average redundancy | 14.2 |  |  |
| Refinement statistics |  |  |  |
| $R$-factor (%) | 22.14 | | |
| $R_{\text{free}}$ (%) | 27.95 | | |
| R.M.S. deviations from ideal values |  |  |  |
| Bond length (Å) | 0.004 |  |  |
| Bond angle (°) | 0.706 |  |  |
| Ramachandran plot (%) |  |  |  |
| Favored | 95.6 |  |  |
| Allowed | 4.19 |  |  |
| No. atoms |  |  |  |
| Total | 24572 |  |  |
| Protein / Water / Ions | 24156 / 404 / 12 |  |  |

## RESULTS

### UPF3B’s domain architecture is homologous to paraspeckle proteins

Due to the clustering of known disease-causing mutations (Figure 1A) within the uncharacterized middle-domain of UPF3B, we computationally analyzed this domain to infer potential structure and function. Quick2D (36) provided a strong consensus indicating several interspersed α-helical regions in the middle-domain (Supplementary Figure S1). Strikingly, two of these stretches are predicted to extend over 70 and 60 residues (P178-P257 comprising the R255K mutation and S328-K390 comprising the R355Q and R366H mutations). DeepCoil and MARCOIL servers (49,50) also indicated that these regions have a >90% probability to form coiled-coil structures (not shown).

Next, we submitted the full-length UPF3B sequence to HHpred to identify related proteins of known structure within the RCSB PDB (51,52). The highest scoring hits comprised RRM domains aligning to the previously solved RRM-L domain of UPF3B (21). Two of the highest scoring hits were for crystal-derived structures of the human splicing factor proline- and glutamine-rich protein (SFPQ) (51) aligning to UPF3B’s RRM-L domain and extending to residue 250 in the middle-domain (Supplementary Figure S2A). SFPQ is a member of the Drosophila Behavior/Human Splicing (DBHS/ paraspeckle) family implicated in subnuclear body formation, transcription, and splicing (53). The DBHS family is structurally characterized by two tandem N-terminal RRM domains, followed by a NonA/paraspeckle domain (NOPS) and a C-terminal coiled-coil (Supplementary Figure S2B,C) (54). The latter, along with the NOPS and RRM domains, facilitates homo- and hetero-dimerization (Supplementary Figure S2B) (55), while UPF3B is monomeric (19). HHpred aligns UPF3B’s RRM-L domain to the second RRM domain of SFPQ (Supplementary Figure S2C) and residues 145-250 of UPF3B’s middle-domain onto the NOPS and coiled-coil regions of SFPQ (Supplementary Figure S2A-C). We generated a SFPQ-based homology model of UPF3B using MODELLER (37) which comprised the RRM-L domain followed by a NOPS-like linker connecting to an α-helical stretch with negative and hydrophobic residues that could facilitate self-association with the RRM-L (Figure 1B). The α-helical region extends into a solvent-exposed, uninterrupted stretch of positively charged residues spanning the length of the modelled helix (Figure 1B).

We produced two UPF3B middle-domain variants expanding over different lengths of the predicted α-helical regions (residues 146-256 and 146-417). Circular dichroism (CD) spectra presented a canonical double dip at 210 nm and 222 nm characteristic for α-helical structure (Figure 1C), with a larger shift in degrees observed for the longer construct. Temperature wavelength scans showed that these constructs unfolded in an unusual non-sigmoidal fashion, indicating gradual unravelling rather than globular domain unfolding, in agreement with the middle-domain adopting an extended α-helical structure (Figure 1C).

### UPF3B middle-domain binds preferentially to double-stranded RNA and displays RNA-induced oligomerization

Members of the DBHS family have been described as “functional aggregators” with the ability to form large aggregates in the presence of DNA (51,54). They act as molecular scaffolds for a large range of different nucleic acids and protein binding partners (51,54). Like DBHS proteins (51), when incubated with single-stranded RNA and DNA oligomers in electrophoretic mobility shift assays (EMSAs), UPF3B displayed multiple binding events leading to the formation of large aggregates unable to migrate into the gel at increased UPF3B concentrations (Supplementary Figure S3A). Affinities for different nucleic acid substrates to UPF3B were determined using fluorescence anisotropy (FA)-based binding assays. The resulting curves show that UPF3B has a ~six-fold higher affinity for single-stranded RNA (ssRNA, 30 nM) compared to ssDNA (190 nM) (Supplementary Figure S3B,C). Intriguingly, UPF3B bound double-stranded (ds) oligomers with considerably higher affinities than their single-stranded counterparts, with dsDNA displaying ~three-fold (61 nM) and dsRNA two-fold (15 nM) higher affinity (Supplementary Figure S3D,E).

With dsRNA being UPF3B’s preferred substrate, we next explored binding to human 40S and 60S ribosomal subunits using surface plasmon resonance (SPR) as these are rich in dsRNA, including double-stranded RNA extension segments (Supplementary Figure S4). In agreement with previous reports of UPF3B binding to both mRNA and ribosomes (19,20), we observed high-affinity interactions with both 40S (38 nM) and 60S (4.3 nM) ribosomal subunits.

### Nucleic acid binding is mediated by UPF3B’s RRM-L and the middle-domain

To determine UPF3B’s minimal RNA-binding domains, stable middle-domain fragments (Figure 2A, Supplementary Figure S5) with and without the RRM-L domain were tested in FA-assays (Figure 2A, Supplementary Figure S6) using a dsRNA oligomer (Supplementary Table S1). The RRM-L domain alone (UPF3B-41-143) displayed very poor affinity for dsRNA (K_D_ of 9.2 μM), corroborating previous EMSA experiments which found no interaction between the RRM-L domain and a ssRNA probe (21). The middle-domain (UPF3B-146-256) revealed only modestly higher affinity (1.8 μM). An extension of this fragment to the complete middle domain (UPF3B-146-417) bound dsRNA with 280 nM affinity, while a construct including the DBHS-homology region with the RRM-L domain and a portion of the middle domain (UPF3B-41-262) was sufficient to bind dsRNA with close to wildtype affinity (37 nM versus 15 nM for UPF3B-WT) (Figure 2A). Further truncation of the middle-domain (UPF3B-41-214) by removal of the positively charged part of the first α-helical region resulted in considerably decreased affinity (1.2 μM) indicating that the first coiled-coil-like domain is essential for UPF3B’s ability to bind dsRNA. Moreover, UPF3B-41-262 retained wildtype-like RNA-induced oligomerization evidenced by multiple band shifts in EMSA (Figure 2B).

**Figure 2.**
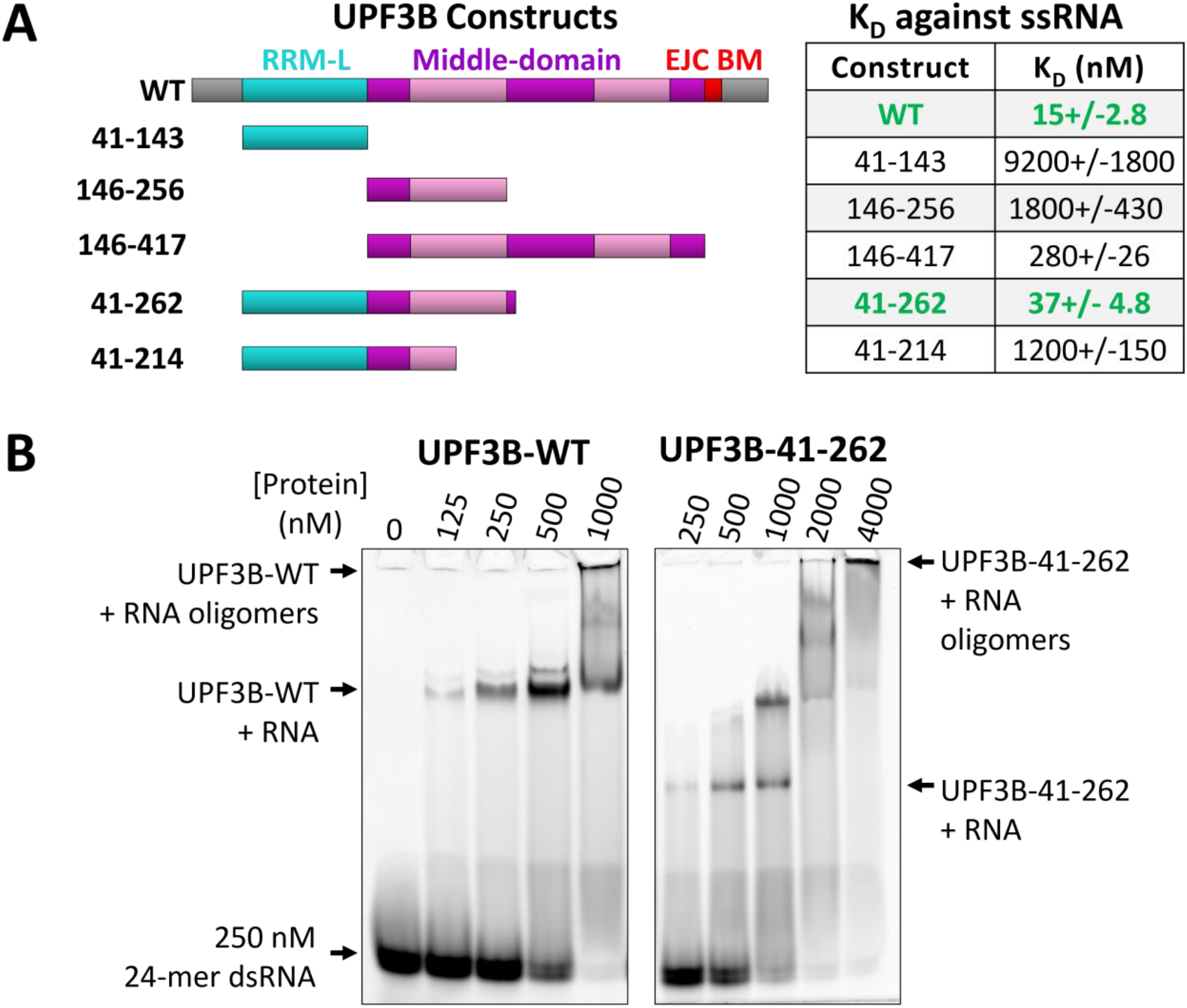
UPF3 RNA binding and RNA-induced aggregation. **(A)** Cartoon representation of the UPF3B constructs (left) utilized for testing dsRNA binding and corresponding dissociation constants determined by fluorescence anisotropy experiments (right) mapping the minimum nucleotide binding interface of UPF3B to its RRM-L and N-terminal part of the middle-domain (residues 41-262, highlighted in green) (binding curves in Supplementary Figure S6). **(B)** Electrophoretic mobility shift assay of UPF3B-WT (left) and UPF3B-41-262 (right) using double-stranded 24-mer RNA (dsRNA) indicating that UPF3B-41-262 retains RNA-induced aggregation behavior.

### Structure of UPF3B in complex with UPF2-MIF4GIII

The interaction between UPF3B’s RRM-L and the UPF2-MIF4GIII domains is well characterized (21) but structural information regarding UPF3B’s middle-domain is lacking. Moreover, it remained unclear if the middle-domain contributes to UPF2 binding. Here, we crystallized a construct comprising UPF3B residues 41-189, comprising the RRM-L and the N-terminal part of middle-domain, in complex with UPF2-MIF4GIII and solved the structure to 2.6 Å resolution (Table 1). In agreement with our UPF3B homology model (Figure 1B), the region after the RRM-L and at the beginning of the middle-domain of UPF3 indeed adopts a NOPS-L domain followed by an α-helix (Figure 3A).

**Figure 3.**
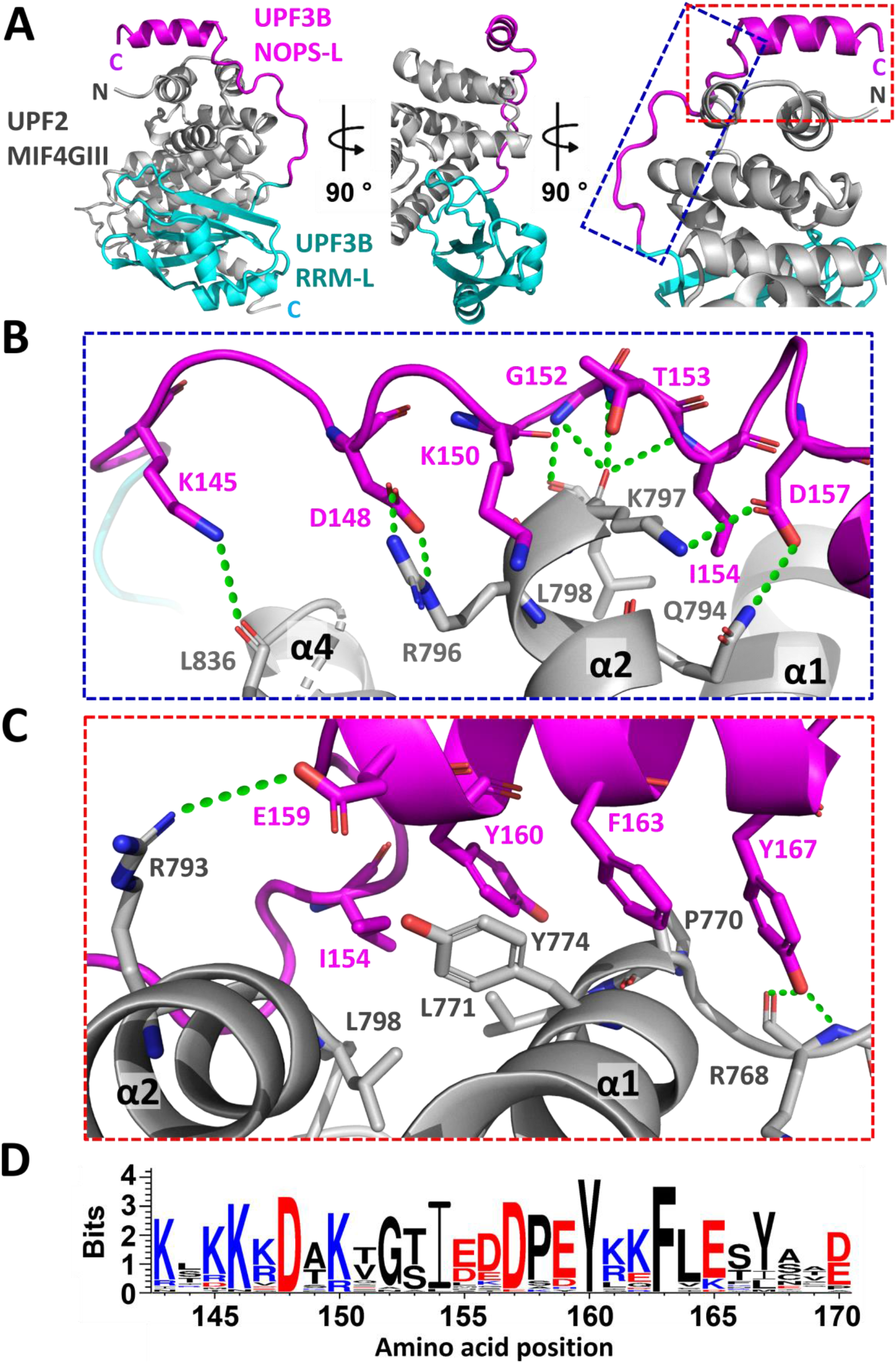
Co-crystal structure of UPF3B-41-189 + UPF2-MIF4GIII complex. **(A)** Cartoon representation of the crystallographic model with three representative views of UPF2-MIF4GIII (grey) complexed with UPF3B-41-189 comprising the RRM-L domain (cyan), the NOPS-L linker and α-helix (magenta). **(B)** Zoomed view (blue box, panel A) showing the residues involved in interactions between the NOPS-L linker with UPF2. **(C)** Zoomed view (red box, panel A) of residues in the α-helical portion contributing to binding of UPF2. Polar and ionic interactions are indicated (green lines). **(D)** WebLogo (38) alignment summarizing sequence conservation across a range of UPF3B homologs (Supplementary Figure S8) for the newly determined region indicating conservation of several of the residues involved in complex formation. The accumulative height of each stack (in bits) indicates the degree of conservation at that position. The height of each individual letter indicates the frequency that amino acid is found at that position.

The RRM-L domain-UPF2 part of the crystal structure aligns very well with a previous UPF3B RRM-L - UPF2-MIF4GIII structure (21) displaying an RMSD of 0.44 Å. In agreement, only minor changes are observed in UPF3B and UPF2-MIF4GIII (Supplementary Figure S7), and the same residues are involved in protein-protein interaction. The following NOPS-L region is solvent-exposed, and conserved NOPS-L residues K145, D148, K150, G152, T153 and D157 form multiple polar and salt-bridge interactions with helices α1, α2, and α4 of UPF2-MIF4GIII (Figure 3B). The subsequent α-helix further contributes to UPF2 interaction, primarily through hydrophobic contacts with helices α1 and α2 of UPF2-MIF4GIII, involving residues I154, Y160, F163 and Y167 (Figure 3C, Figure 4A,B). Moreover, UPF3B-E159 at the beginning of the α-helical region forms a salt-bridge with R793 of UPF2 (Figure 3C). Alignments of UPF3B and UPF3A show that this region (residues 143-170) is highly conserved in plants and animals, with particularly strong conservation for residues D148, I154, D157, Y160, and F163 (Figure 3D, Supplementary Figure S8). Together, the RRM-L, NOPS-L and α-helix of UPF3B wrap around UPF2-MIF4GIII leading to an intimate interaction.

**Figure 4.**
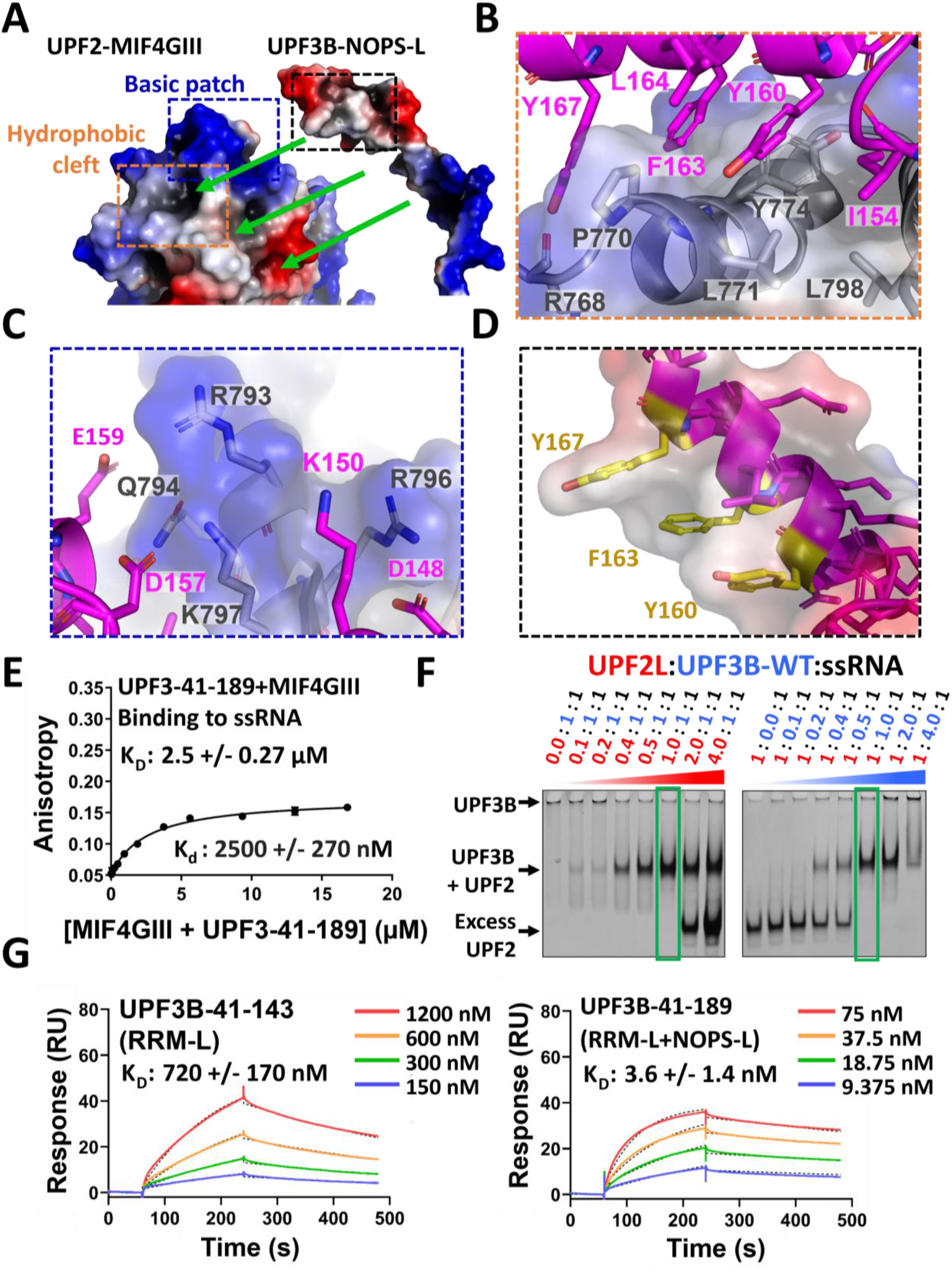
UPF3B-UPF2 interface analysis. **(A)** Electrostatic surface potential of UPF2-MIF4GIII and the NOPS-L domain of UPF3B. Blue, red and white surface indicates positive, negative, and hydrophobic surface potential, respectively. **(B)** Zoom in of hydrophobic cleft (orange box, panel A) interactions between UPF3B-NOPS-L (magenta cartoon) and UPF2-MIF4GIII (electrostatic surface and grey cartoon). **(C)** Zoom in of UPF2-MIF4GIII basic patch residues (blue box, panel A) and their interactions with UPF3B-NOPS-L. **(D)** Zoom in of UPF3B-NOPS-L α-helical stretch (black box, panel A) residues involved in hydrophobic interaction with UPF2-MIF4GIII hydrophobic cleft (including disease-causing mutant Y160). Aromatic residues selected for mutagenesis are highlighted in yellow. **(E)** Fluorescence anisotropy binding curves of the complex co-crystallized in this study (UPF3B-41-189 + UPF2-MIF4GIII) titrated against HEX-labelled 24mer ssRNA indicating a strong inhibition of UPF2-RNA binding in the presence of UPF3B-41-189. Protein titrations were carried out in triplicate and error bars plotted via standard deviation before fitting a single component binding equation in GraphPad Prism to calculate KD values. **(F)** Coomassie-stained native 4-20% Novex gels loaded with ssRNA+UPF3B-WT incubated with increasing amounts of UPF2L (left) and with ssRNA+UPF2L incubated with increasing amounts of UPF3B-WT (right). Green boxes highlight UPF3B:UPF2 complexes at 1:1 stoichiometry. **(G)** SPR sensorgrams of serial dilution injections (colored lines) of RRM-L (UPF3B-41-143, left) and RRM-L+NOPS-L (UPF3B-41-189, right) over biotinylated-UPF2-MIF4GIII immobilized on a streptavidin-coated chip, highlighting a 200-fold affinity change. Sensorgrams were globally fitted with a 1:1 binding model within the T200 Biacore Evaluation Software (black dotted lines) to determine indicated dissociation constants.

### UPF3B interferes with RNA binding by UPF2-MIF4GIII

Conserved acidic residues D148, D157, and E159 of UPF3B form ionic interactions with a basic patch of UPF2-MIF4GIII comprising residues R793, R796, K797 and Q794 (Figure 4A,C). This basic patch of UPF2 was previously shown to bind RNA *in vitro* (21). Considering that these residues coordinate UPF3B in our structure, we postulated that UPF2-MIF4GIII RNA binding would be impaired when in complex with UPF3B-41-189. In agreement, we observed a 4-fold decrease in UPF2-MIF4GIII affinity for ssRNA (2.5 μM) in the presence of UPF3B-41-189 compared to UPF3B RRM-L alone (0.62 μM) or in the absence of UPF3B (0.7 μM) in FA assays (Figure 4E, Supplementary Figure S6F,G). We conclude that the UPF3B’s NOPS-L domain outcompetes ssRNA for UPF2-MIF4GIII binding.

### UPF2 interferes with RNA-induced oligomerization of UPF3B

We next investigated RNA and UPF3B complex formation using a larger UPF2 construct with wildtype activity (UPF2L, amino acids 120-1227) (15,16). UPF3B-WT was incubated with ssRNA concentrations that lead to aggregate formation in EMSAs (Figure 4F, Supplementary Figure S3A left). Subsequent addition of equimolar amounts of UPF2L dissolved the large aggregates and led to formation of UPF3B-UPF2L complexes with defined stoichiometry (Figure 4F, green box). Inversely, titration of UPF3B-WT into a fixed concentration of ssRNA and UPF2L leads to formation of defined UPF2L-UPF3B complexes and at higher concentrations to formation of aggregates which sequester UPF2L (Figure 4F). Together, this data suggests that excess UPF2 alters the RNA-binding behavior of UPF3B and inhibits the formation of higher-order oligomers.

### UPF3B-Y160 is critical for high-affinity UPF2 binding

We further explored the contribution of the NOPS-L and the following α-helix to the UPF3B-UPF2 interface. Using SPR, we determined a K_D_ for UPF3B-41-189 of 3.6 nM, which is 200-fold lower than the RRM-L alone (720 nM) (Figure 4G). Next, we investigated the contribution of conserved residues Y160, F163, and Y167, which are buried in a hydrophobic cleft formed by UPF2-MIF4GIII residues P770, L771, Y774, L798 (Figure 4A,B,D). Importantly, Y160D is implicated in XLID disease phenotypes, but the molecular basis of this remained enigmatic (25,26,28). Moreover, UPF3B residues Y160 and Y167 are phosphorylated when expressed in insect cells (Supplementary Figure S9) and in human cells (56,57). We tested UPF2-MIF4GIII binding of UPF3B-41-189 mutants Y160A, Y160D (phosphomimetic/ disease mutation), F163A, Y167A, Y167D (phosphomimetic), and double mutants Y160A+Y167A and Y160D+Y167D (Figure 5A, Supplementary Figure S10). UPF3B’s Y160A and Y160D mutants reduced the affinity to 88 nM (24-fold reduction) and to 140 nM (39-fold reduction) respectively, compared to 3.6 nM for UPF3B-1-189. Similarly, the F163A mutation led to a K_D_ of 90 nM (25-fold reduction). In contrast, UPF3B mutations Y167A and Y167D had close to no impact, resulting in K_D_’s of 3.3 nM and 7.4 nM for UPF2-MIF4GIII respectively. In agreement, UPF3B double mutants behaved like single Y160 mutants with both Y160A/Y167A and Y160D/Y167D having K_D_’s of ~130 nM. In summary, phosphorylation of tyrosine residue 160 or mutation to aspartate, both result in reduced affinities by preventing accommodation of this residue into the hydrophobic cleft of UPF2-MIF4GIII.

**Figure 5.**
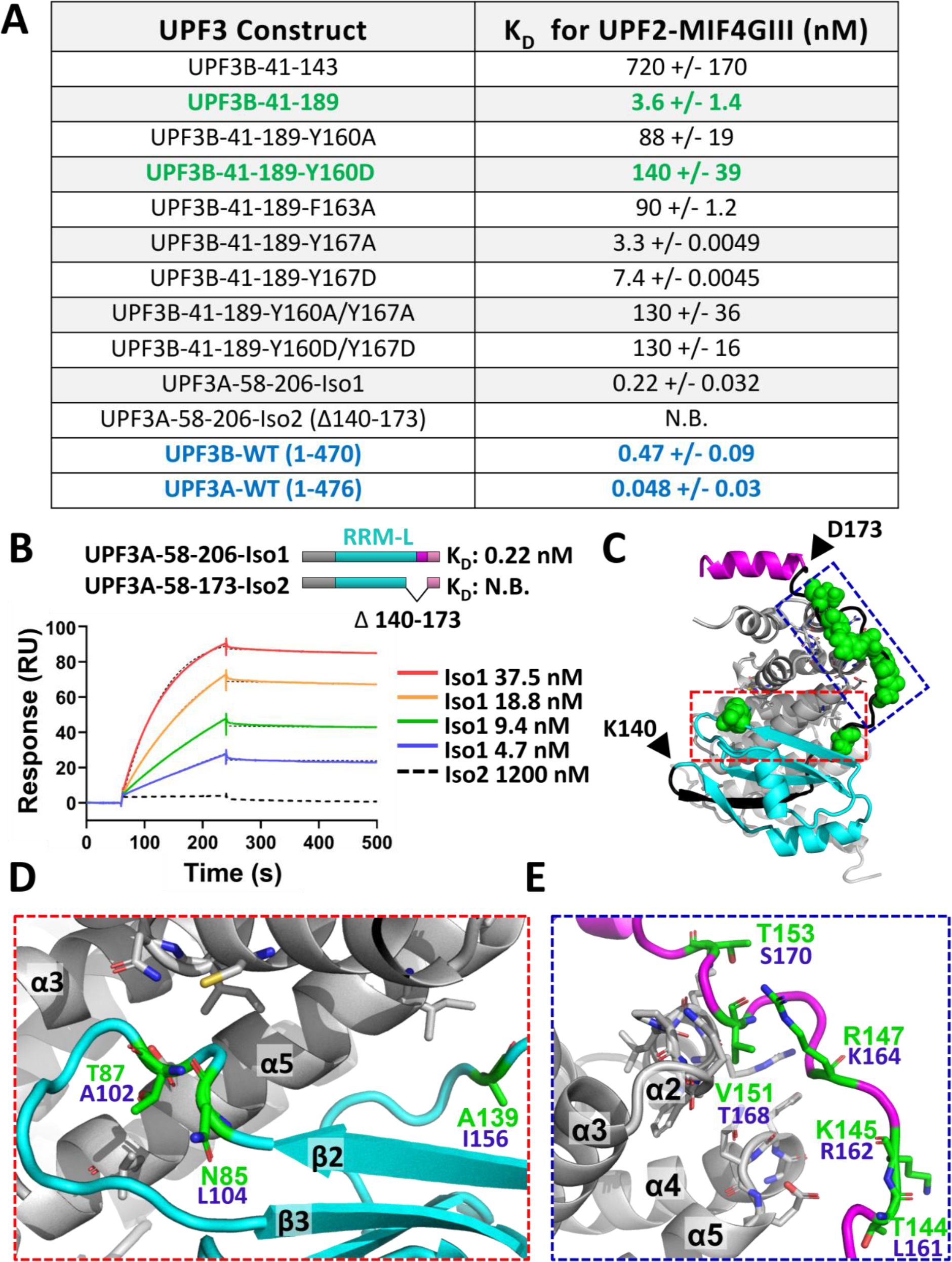
Characterization of interactions of UPF3B variants and UPF3A with UPF2-MIF4GIII. **(A)** Table summarizing the KD values determined in this study by SPR between immobilized UPF2-MIF4GIII and UPF3 constructs, highlighting a 40-fold affinity decrease due to Y160D mutation (green) and a 10-fold higher affinity of UPF3A compared to UPF3B (blue). Binding curves are shown in Supplementary Figures S10,S11. N.B. stands for no binding. **(B)** Schematic of UPF3A-58-206-Iso1 and UPF3A-58-173-Iso2 constructs and representative SPR sensorgrams produced via injections of these proteins over immobilized UPF2-MIF4GIII. **(C)** The co-crystal structure of UPF3B-41-189 + UPF2-MIF4GIII derived in this study highlighting residues found at the UPF2-UPF3B interface that are non-conserved in UPF3A (green spheres). The UPF3A-isoform2 deletion region is colored black. Start (K140) and end (D173) of the deletion is highlighted with black triangles. **(D)** Zoom in of RRM-L residues at the UPF2 interface that are non-conserved in UPF3A (red box, panel C). Residue label numbers in green are for UPF3B and blue for UPF3A. **(E)** Zoom in of UPF3B NOPS-L residues at the UPF2 interface that are non-conserved in UPF3A (blue box, panel C).

### Impact of UPF3A isoforms on UPF2 binding

The UPF3B Y160D mutation leads to an upregulation of UPF3A protein levels in affected families (25,28,32). Two splice variants of UPF3A exist in humans; isoform1 which retains exon4 and isoform2 which excludes exon 4 (Δ141-173) thereby deleting the β5 strand of the RRM-L and the entire NOPS-L domain (Figure 5B,C). We generated corresponding UPF3A constructs with the same boundaries as UPF3B-41-189 (UPF3A-58-206-Iso1, UPF3A-58-173-Iso2) comprising the RRM-L and N-terminal part of the middle-domain. Surprisingly, UPF3A-58-206-Iso1 had an affinity of 220 pM to UPF2-MIF4GIII, binding 16-fold tighter than UPF3B-41-189 (Figure 5A,B). In contrast, UPF3A-58-173-Iso2 showed no measurable response in SPR assays (Figure 5A,B), explainable by the deletion of a large part of the UPF2-interaction surface (Figure 5C). In agreement, previous studies find that unlike UPF3A-isoform1, isoform2 is inactive in NMD and unable to pull-down UPF2 in co-immunoprecipitation experiments (33,58). To corroborate this finding, we next used full-length UPF3A-WT and UPF3B-WT in SPR analyses (Supplementary Figure S11) and determined affinities of 470 pM for UPF3B-WT and 48 pM for UPF3A-WT for UPF2-MIF4GIII, indicating a similar higher affinity (~10-fold) for UPF3A (Figure 5A).

Scrutinizing our UPF3B and UPF2-MIF4GIII complex structure, we identified eight UPF3A-isoform1 residues which are not conserved in UPF3B and at the interface of UPF2-MIF4GIII: three within RRM-L (Figure 5C,D) and five in NOPS-L (Figure 5C,E). UPF3B N85 and T87 found within a hairpin between β2 and β3 of the RRM-L correspond to A102 and L104 in UPF3A which could promote hydrophobic interactions with MIF4GIII helices α3 and α5. In addition, UPF3B A139 substitution to UPF3A I156 located at the C-terminus of RRM-L could promote hydrophobic interactions with helix α5 of UPF2-MIF4GIII. In the NOPS-L, we predict that K145 to R162 as well as V151 to T168 will add further polar interactions with UPF2-MIF4GIII residues on the loops connecting α2-α3 and α4-α5, leading to an even more intimate interface. In summary, UPF3A binds with higher affinity and thus could outcompete UPF3B in a direct competition for UPF2 binding.

## DISCUSSION

We here describe the architecture of UPF3B, show that UPF3B shares structural and functional homology with paraspeckle/DBHS proteins, and identify a novel interface between UPF2 and UPF3B highlighting the importance of UPF3B’s middle-domain and of residue Y160. Our findings dissect two known key roles of UPF3B in NMD:

1. UPF3B binds mRNA and ribosomal subunits (19,20). *In vitro*, UPF3B slows down translation termination and supports ribosome dissociation after peptide release (19). Consistent with its homology to paraspeckle proteins, we show that UPF3B undergoes RNA-induced oligomerization at high concentrations reliant on the N-terminal part of the middle-domain (Figure 2B). However, UPF3B is a nucleo-cytoplasmic shuttling protein showing enrichment in nucleoli (25), while localization in nuclear or cytoplasmic granules is not reported so far. Thus, it is unclear if cellular concentrations of UPF3B suffice to aggregate RNA, and the functional consequences remain unexplored. UPF3B-RNA cross-linking studies showed that UPF3B interacts with mRNA 15-30 nucleotides upstream of exon-exon junctions (20) indicating that UPF3B’s mRNA binding may assist in positioning the NMD machinery. Interestingly, mRNA binding of UPF3A recently has been implicated in the genetic compensation response (59).
2. The UPF2-UPF3B complex is essential for activation of UPF1 helicase (15,16) and of SMG1 kinase which phosphorylates UPF1 leading to recruitment of nucleases and mRNA decay (17,18). We show that UPF2-UPF3B complex formation alters the RNA binding of UPF3B and interferes with RNA-induced UPF3B oligomerization. The intimate UPF2-UPF3B interface involves the NOPS-L linker and the first α-helix of the middle-domain in addition to the RRM-L domain of UPF3B (21). Our structure reveals how the disease-causing Y160D mutation in UPF3B located in the NOPS-L α-helix impairs binding to a hydrophobic cleft in UPF2-MIF4GIII, reducing the binding affinity by ~40-fold. In cells, weakening the UPF2-UPF3B interaction through UPF3B mutations in the RRM-L leads to upregulation of UPF3A levels and NMD inhibition (32). Importantly, UPF3A upregulation appears to be achieved through stabilization of the inherently unstable free UFP3A protein via UPF2 binding (32). Surprisingly, we found that UPF3A had a significantly higher affinity (~10-fold) for UPF2 than UPF3B (Figure 5A) which raises the question how UPF3B can outcompete UPF3A for UPF2 binding. Cells could achieve this through differences in tissue-dependent ratios of UPF3B and UPF3A, tissue-dependent splicing of UPF3A (24,32,34), or via phosphorylation of UPF3A and UPF3B at Y177 and Y160 respectively and then exploit resulting changes in UPF2 binding affinity to finetune NMD efficiency.

## DATA AVAILABILITY

Coordinate and structure files have been deposited to the Protein Data Bank with accession code 7NWU. All other data are available by request.

## SUPPLEMENTARY DATA

Supplementary data is available at NAR online.

## ACKNOWLEDGEMENTS

We thank all members of the Berger and Schaffitzel teams for their assistance and advice as well as Gabriele Neu-Yilik and Imre Berger for critical reading of the manuscript. We acknowledge the staff of the BrisSynBio BioSuite including Peter Wilson and the beamline scientists at the Diamond Light Source beamline I04-1 (via BAG access) for their assistance with crystallographic studies.

## AUTHOR CONTRIBUTIONS

C.S., J.C.B., and K.T.P. designed the study. J.C.B, K.T.P., and C.S. wrote the manuscript. J.C.B. conducted the structural predictions and homology modelling. J.C.B. and K.T.P. cloned and expressed the constructs used throughout the study. J.C.B. conducted the circular dichroism spectroscopy. J.C.B. and J.A.S. conducted the electrophoretic mobility-shift assays. J.C.B. and J.A.S. conducted the fluorescence anisotropy assays. K.T.P. and J.C.B. conducted the crystallographic screening, data collection, and model refinement. J.C.B. generated the sequence conservation plots. J.C.B. and K.T.P. conducted the surface plasmon resonance assays.

## COMPETETING INTERESTS

Authors declare that they have no competing interests.

