## Supplementary Data for "Molecular basis of neurodevelopmental disorder-causing mutation in nonsense-mediated mRNA decay factor UPF3B"

**This PDF file includes:**

Supplementary Figures S1 to S11

Supplementary Tables S1 and S2

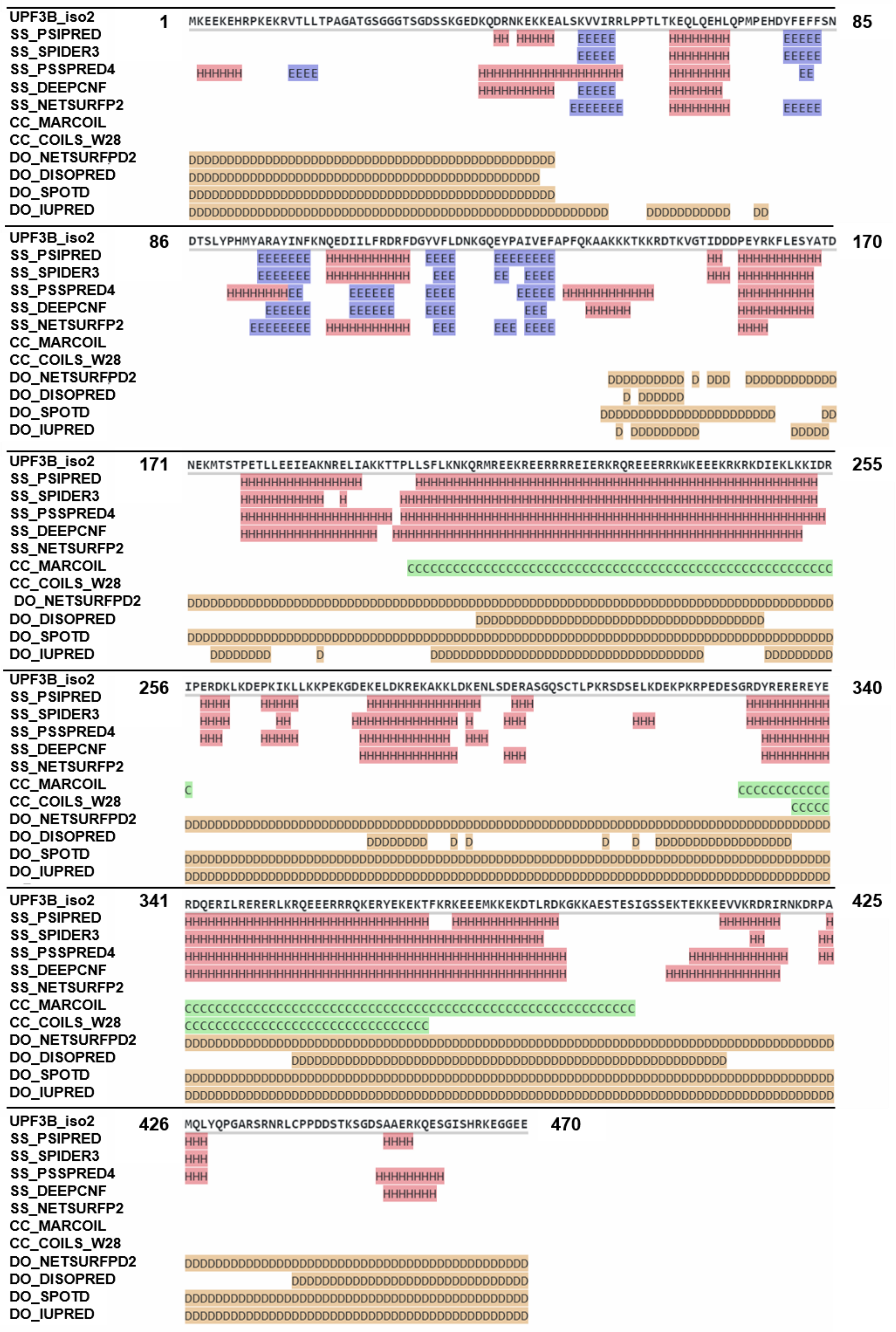
UPF3B_iso2 1 MKEEKEHRPKEKRVTLLTPAGATGSGGGTSGDSSKGEDKQDRNKEKKEALSKVVIRRLPPTLTKEQLQEHLQPMPEHDYFEFFSN 85
UPF3B_iso2 86 DTSLYPHMYARAYINFKNQEDIILFRDRFDGYVFLDNKGQEYPAIVEFAPFQKAAKKKTKKRDTKVGTIDDDPEYRKFLESYATD 170
UPF3B_iso2 171 NEKMTSTPETLLEEIEAKNRELIAKKTTPLLSFLKNKQRMREEKREERRRREIERKRQREEERRKWKEEEKRKRKDIEKLKKIDR 255
UPF3B_iso2 256 IPERDKLKDEPKIKLLKKPEKGDEKELDKREKAKKLDKENLSDERASGQSCTLPKRSDSELKDEKPKRPEDESGRDYREREREYE 340
UPF3B_iso2 341 RDQERILRERERLKRQEEERRRQKERYEKEKTFKRKEEEMKKEKDTLRDKGKKAESTESIGSSEKTEKKEEVVKRDRIRNKDRPA 425
UPF3B_iso2 426 MQLYQPGARSRNRLCPPDDSTKSGDSAAERKQESGISHRKEGGEE 470
SS_PSIPRED
SS_SPIDER3
SS_PSSPRED4
SS_DEEPCNF
SS_NETSURFP2
CC_MARCOIL
CC_COILS_W28
DO_NETSURFPD2
DO_DISOPRED
DO_SPOTD
DO_IUPRED

**Supplementary Figure S1. UPF3B secondary structure prediction.** Quick2D prediction (Max Planck Institute Bioinformatics Tool Kit) of full-length wildtype UPF3B (isoform2), covering several different secondary structure prediction servers (36). An overview of predicted secondary structure features including $\alpha$-helices (red, H), b-sheets (blue E), coiled coils (green C) and disordered regions (brown D) is shown for each server underneath the primary protein sequence.

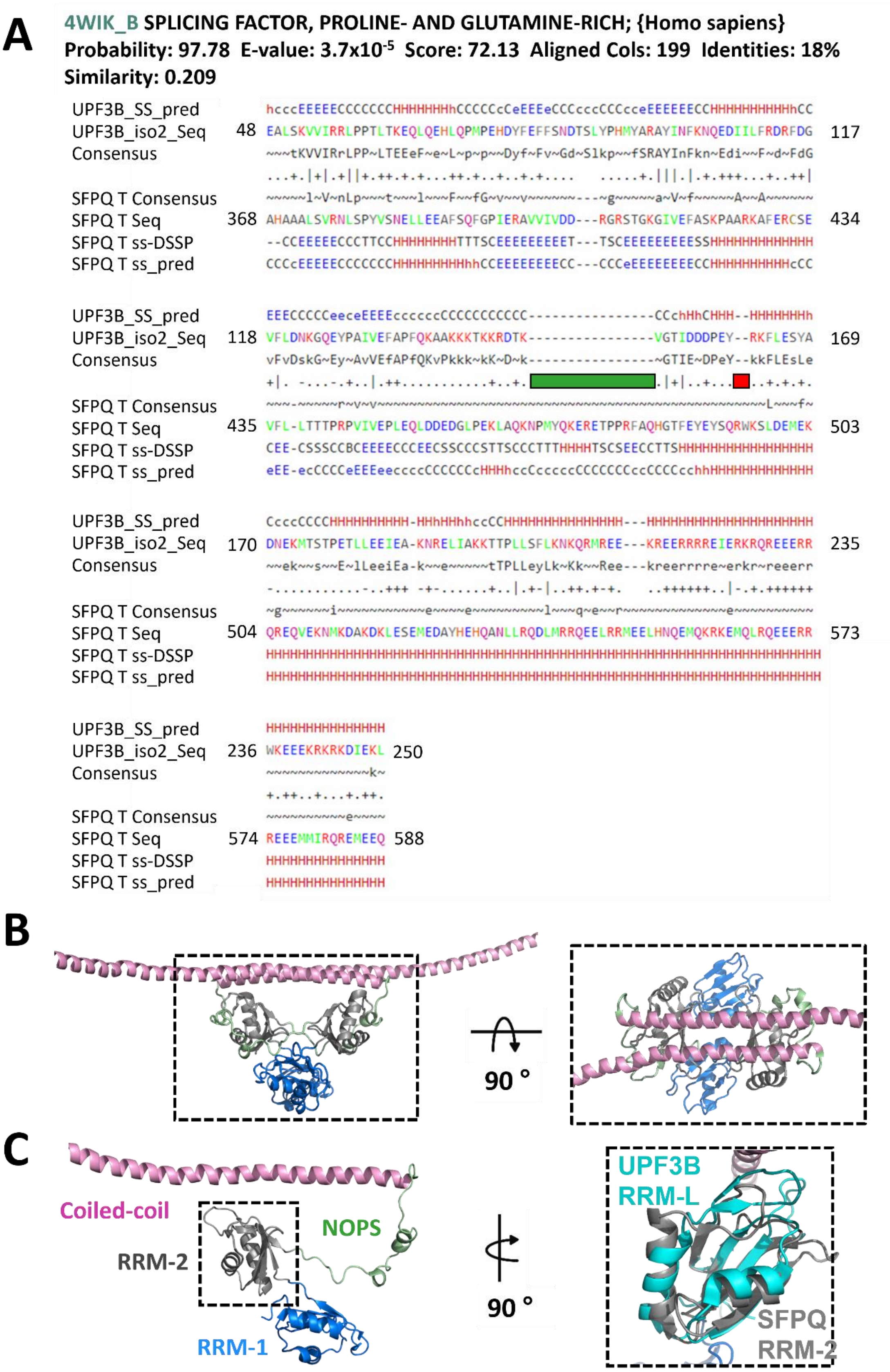
A
4WIK_B SPLICING FACTOR, PROLINE- AND GLUTAMINE-RICH; {Homo sapiens}
Probability: 97.78 E-value: 3.7x10-5 Score: 72.13 Aligned Cols: 199 Identities: 18%
Similarity: 0.209
UPF3B_SS_pred
UPF3B_iso2_Seq
Consensus
SFPQ T Consensus
SFPQ T Seq
SFPQ T ss-DSSP
SFPQ T ss_pred
B
90 °
C
Coiled-coil
RRM-2
NOPS
RRM-1
90 °
UPF3B
RRM-L
SFPQ
RRM-2

**Supplementary Figure S2. SFPQ HHpred homology detection hit for UPF3B. (A)** UPF3B Isoform2 HHpred homology detection hit alignment (36) for PDB ID 4WIK (SFPQ-369-598 homodimer) (51). The probability value describes the likelihood of the hit having homology to the query. The E-value describes how many hits with a better probability score would be expected if the database contained only unrelated hits (36). Homology between the sequences is indicated by the matching of the consensus sequences and by the presence of conserved motifs marked by '+' and '|' signs between alignments. Green and red bars denote the locations of the UPF3B "deletions" in proximity to SFPQ residues Y490 and W494, which are implicated in dimerization of DBHS family members (46). **(B)** Side (left) and top (right) views of the crystal structure of homodimeric SFPQ-276–598 (PDB ID: 4WIJ) (51). **(C)** Left: Monomer of SFPQ-276–598 highlighting DBHS family core domains. The N-terminal tandem RRM1-RRM2 folds are depicted in blue and grey, respectively. The NOPS domain is colored in light green and the antiparallel coiled-coil domain in pink. Right: Superimposition of UPF3B's RRM-L domain (1UW4, cyan) (21) onto the RRM-2 domain of SFPQ indicating a high degree of homology.

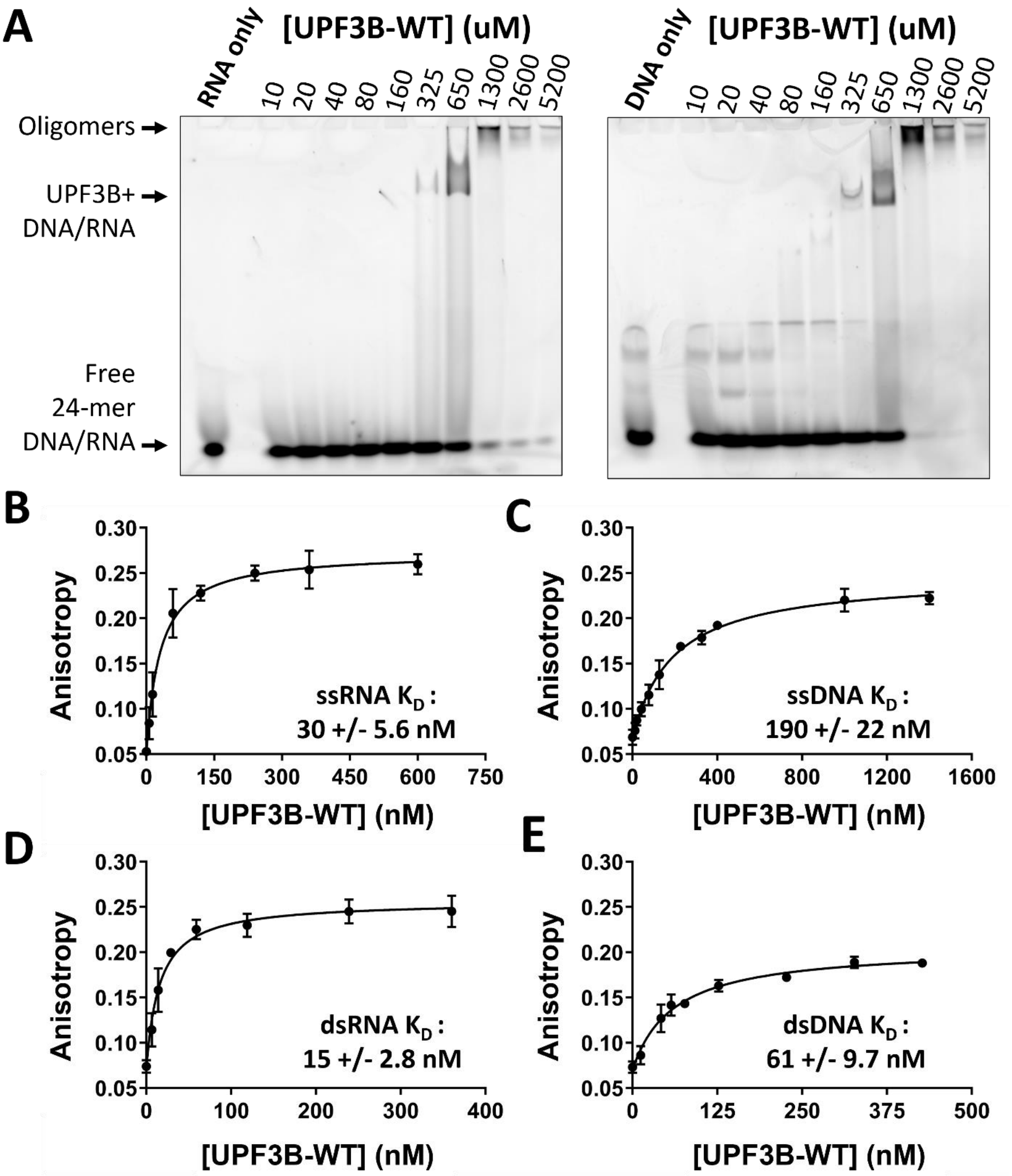
A
RNA only
[UPF3B-WT] (uM)
10 20 40 80 160 325 650 1300 2600 5200
DNA only
[UPF3B-WT] (uM)
10 20 40 80 160 325 650 1300 2600 5200
Oligomers
UPF3B+ DNA/RNA
Free 24-mer DNA/RNA
B
Anisotropy
[UPF3B-WT] (nM)
ssRNA $K_D$ : 30 +/- 5.6 nM
C
Anisotropy
[UPF3B-WT] (nM)
ssDNA $K_D$ : 190 +/- 22 nM
D
Anisotropy
[UPF3B-WT] (nM)
dsRNA $K_D$ : 15 +/- 2.8 nM
E
Anisotropy
[UPF3B-WT] (nM)
dsDNA $K_D$ : 61 +/- 9.7 nM

**Supplementary Figure S3. UPF3B nucleic acid-induced aggregation and substrate preference. (A)** EMSAs of UPF3B-WT with single-stranded 24-mer RNA (left) and single-stranded 24-mer DNA (right) indicating aggregation behavior of UPF3B in the presence of both ssDNA and ssRNA. Representative gel image shown of independent triplicate EMSAs conducted. **(B-E)** Fluorescence anisotropy binding curves for UPF3B-WT with 24-mer ssRNA **(B)**, ssDNA **(C)**, dsRNA **(D)** and dsDNA **(E)** indicating a preference for RNA over DNA, as well as double-stranded oligonucleotides over single-stranded oligonucleotides. Protein titrations and measurements were carried out in triplicate to produce error bars by standard deviation before fitting a single component binding equation (see Methods) in GraphPad Prism to calculate the reported dissociation constants highlighted in each graph. Sequences of the oligomers are listed in Supplementary Table S1.

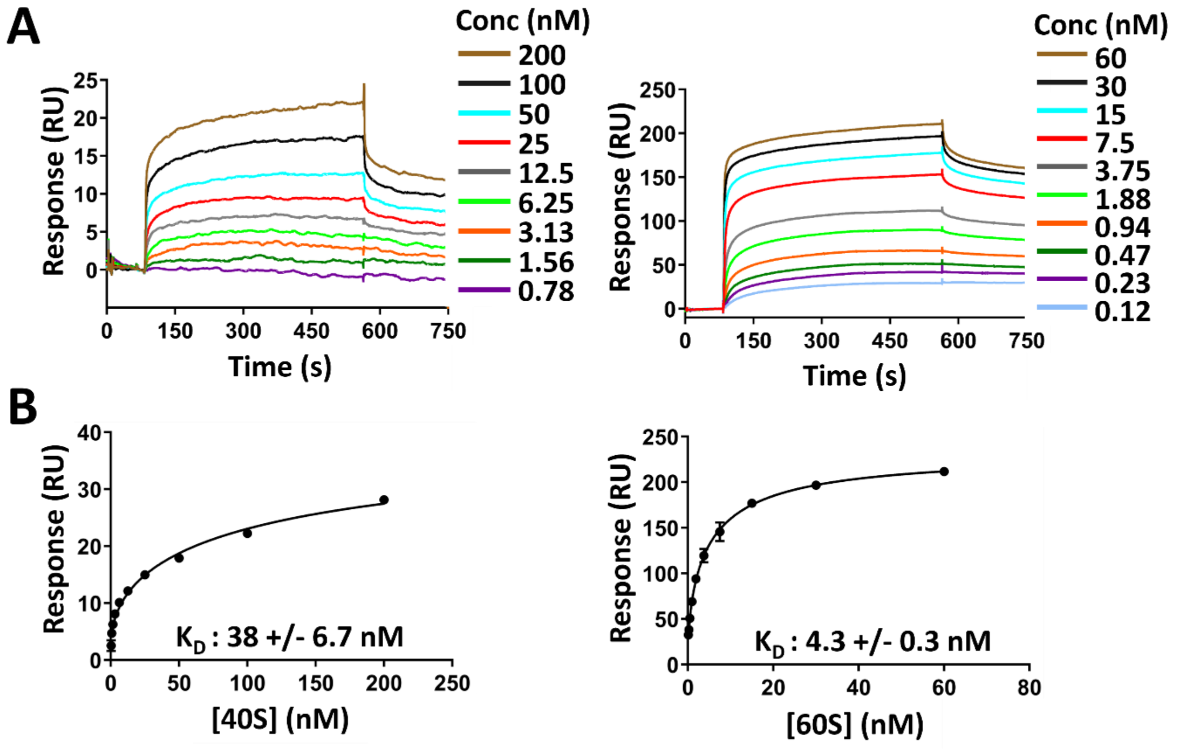


**Supplementary Figure S4. Human 40S and 60S ribosomal subunit binding of UPF3B. (A)** Representative sensorgrams for each analyte concentration of 40S (left) and 60S (right) subunits injected over immobilized biotinylated Avi-UPF3B-WT. **(B)** Steady-state fit plots generated by plotting responses produced by the small ribosomal subunit 40S (left) and the large ribosomal subunit 60S (right) at 450 seconds post injection vs concentration of analyte. Data points were fitted with the single component binding equation in GraphPad Prism to calculate the reported $K_D$ values.

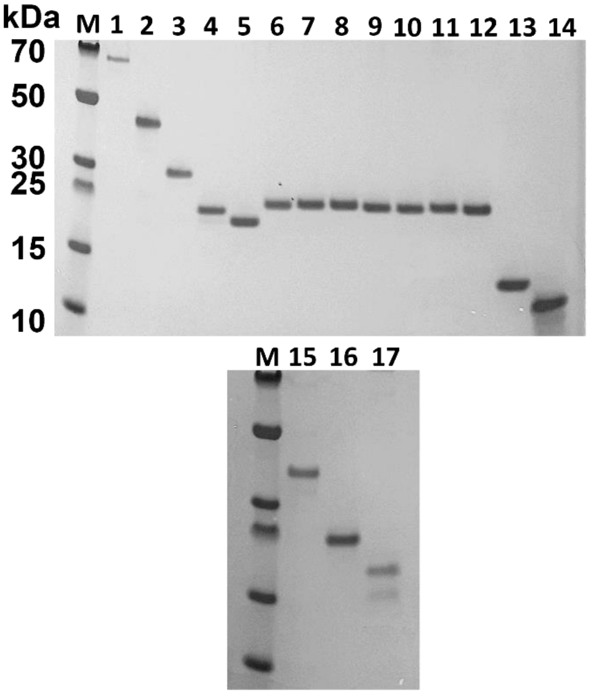


| Lane | Construct |
| --- | --- |
| 1 | UPF3B-WT (1-470) |
| 2 | UPF3B-146-417 |
| 3 | UPF3B-41-262 |
| 4 | UPF3B-41-214 |
| 5 | UPF3B-41-189 (TEV cleaved) |
| 6 | UPF3B-41-189-Y160A |
| 7 | UPF3B-41-189-Y160D |
| 8 | UPF3B-41-189-F163A |
| 9 | UPF3B-41-189-Y167A |
| 10 | UPF3B-41-189-Y167D |
| 11 | UPF3B-41-189-Y160A/Y167A |
| 12 | UPF3B-41-189-Y160D/Y167D |
| 13 | UPF3B-41-143 |
| 14 | UPF3B-146-256 |
| 15 | UPF2-MIF4GIII (761-1054) |
| 16 | UPF3A-58-206-Iso1 |
| 17 | UPF3A-58-173-Iso2 |

**Supplementary Figure S5. SDS-PAGE of purified protein samples used in this study.** Coomassie-stained 4-12% NuPAGE Bis-Tris SDS-PAGE gels of the protein samples produced in this study loaded at ~0.5 µg. M stands for molecular weight marker.

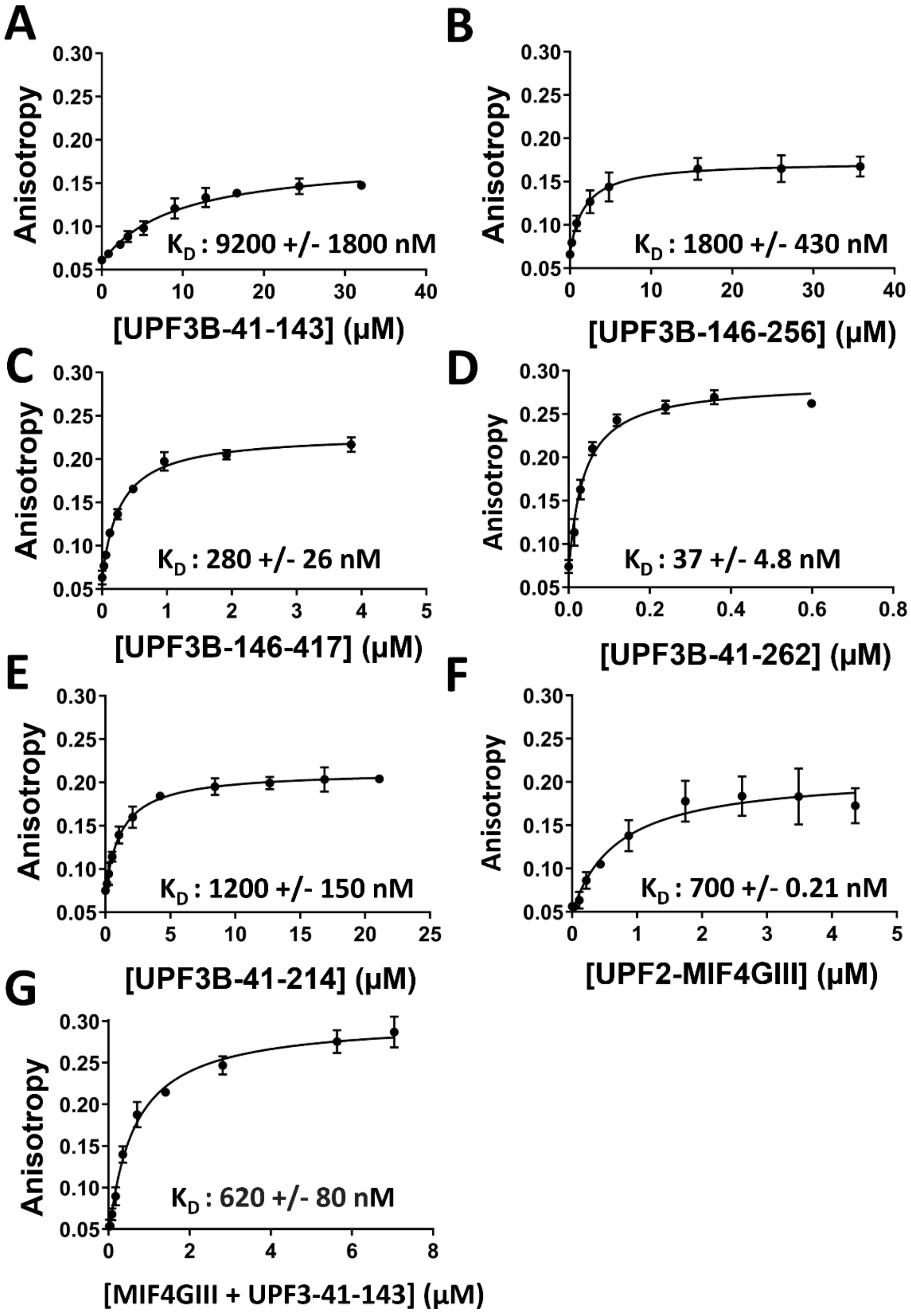
A
Anisotropy
K$_D$ : 9200 +/- 1800 nM
[UPF3B-41-143] (μM)
B
K$_D$ : 1800 +/- 430 nM
[UPF3B-146-256] (μM)
C
K$_D$ : 280 +/- 26 nM
[UPF3B-146-417] (μM)
D
K$_D$ : 37 +/- 4.8 nM
[UPF3B-41-262] (μM)
E
K$_D$ : 1200 +/- 150 nM
[UPF3B-41-214] (μM)
F
K$_D$ : 700 +/- 0.21 nM
[UPF2-MIF4GIII] (μM)
G
K$_D$ : 620 +/- 80 nM
[MIF4GIII + UPF3-41-143] (μM)

**Supplementary Figure S6. RNA binding of UPF3B variants and of UPF2-MIF4GIII. (A-E)** Fluorescence anisotropy binding curves for UPF3B-41-143 **(A)**, UPF3B-146-256 **(B)**, UPF3B-146-417 **(C)**, UPF3B-41-262 **(D)**, and UPF3B-41-214 **(E)** titrated against 24mer HEX-labelled dsRNA. **(F-G)** Binding curves for UPF2-MIF4GIII **(F)** and UPF2-MIF4GIII complexed with UPF3B-41-143 / the RRM-L domain **(G)** vs 24mer HEX-labelled ssRNA. Protein titrations were carried out in triplicate and error bars plotted via standard deviation before fitting using the single component binding equation in GraphPad Prism to calculate the reported dissociation constants.

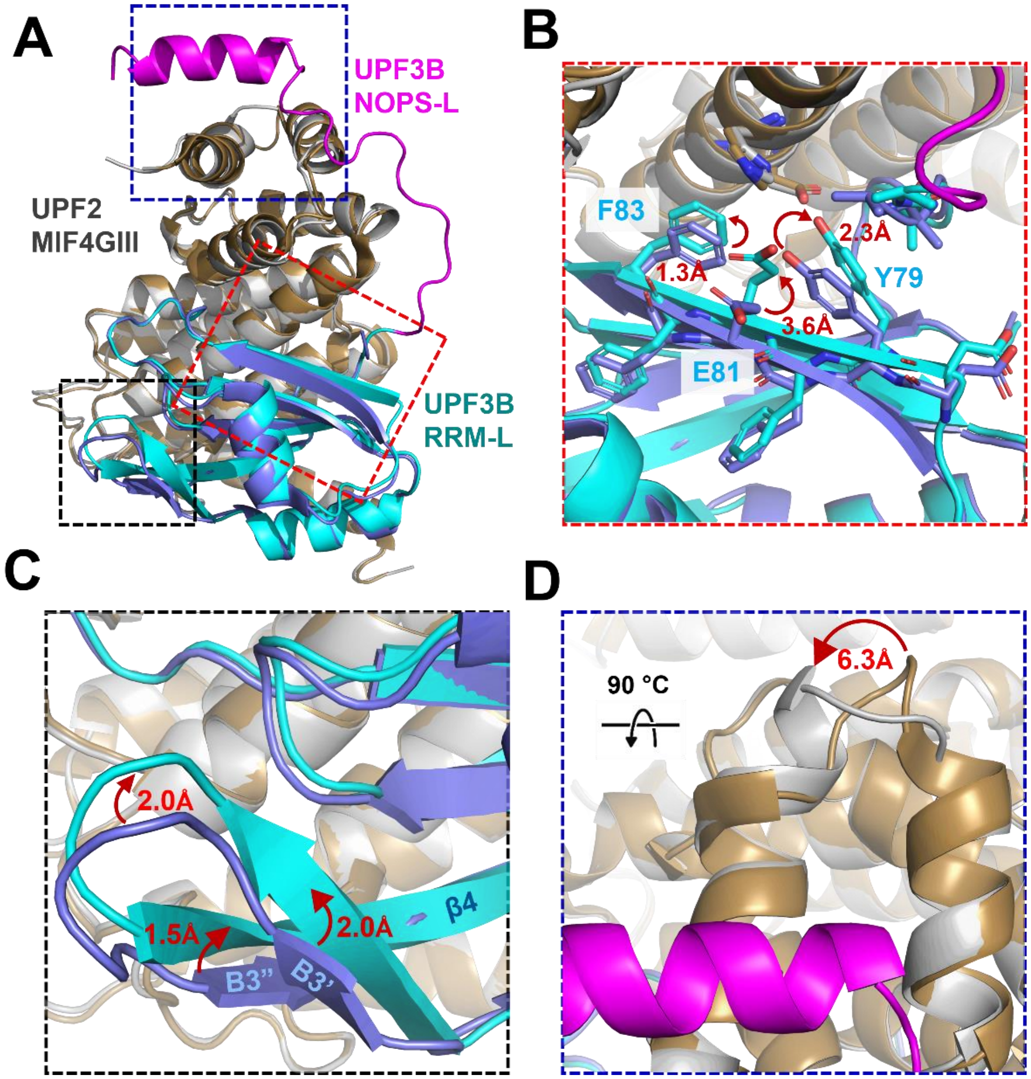


**Supplementary Figure S7. Comparison of co-crystal structures reported in this study and of the complex between UPF3B RRM-L and UPF2-MIF4GIII** (PDB ID 1UW4) (21)**.** UPF3B is colored cyan (RRM-L) and magenta (NOPS-L) (our structure), and light blue (RRM-L, PDB ID 1UW4); UPF2-MIF4GIII is grey (our structure) and gold (PDB ID 1UW4). **(A)** Alignment of the two crystal structures. **(B)** Zoom in on the interface region comprising the β2-strand of UPF3B (highlighted with a red box in panel A). **(C)** Zoom in on B3’, B3”, and β4 (1UW4) and continuous sheets B3’/3” and β4 (our structure) of UPF3B (highlighted with a black box in panel A). **(D)** Zoom in on helices α-1 and α-2 of UPF2 (highlighted with a blue box in panel A).

```
Homo-sapiens-UPF3B                      KTKKRDTKVGTIDDDPEYRKFLESYATD
Pan-troglodytes-UPF3B                   KTKKRDTKVGTIDDDPEYRKFLESYATD
Macaca-mulatta-UPF3B                    KTKKRDTKVGTIDDDPEYRKFLESYATD
Bos-taurus-UPF3B                        KTKKRDTKVGTIDDDPEYRKFLESYAAD
Canis-lupus-familiaris-UPF3B            KTKKRDTKVGTIDDDPEYRKFLESYAAD
Rattus-norvegicus-UPF3B                 KIKKRDTKVGTIEDDPEYRKFLESYATD
Mus-musculus-UPF3B                      KIKKRDTKVGTIEDDPEYRKFLESYATD
Tetraodon-nigroviridis-UPF3B            RSKKRDAKCGTINEDPEYKKFLEFYNGD
Danio-rerio-UPF3                        RSKKKDAKSGTIDDDADYKKFLEFYNGD
Taeniopygia-guttata-UPF3                KSKKKDAKTGTIEDDPEYKKFLESYSAD
Gallus-gallus-UPF3                      KSKKKDAKTGTIEDDPEYKKFLESYSAD
Xenopus-laevis-UPF3                     KSKKQDSKIGTIDEDPEYRKFLDSYTMD
Homo-sapiens-UPF3A                      KLRKKDAKTGSIEDDPEYKKFLETYCVE
Pan-troglodytes-UPF3A                   KLRKKDAKTGSIEDDPEYKKFLETYWVE
Macaca-mulatta-UPF3A                    KLKKKDAKTGSIEDDPEYKKFLETYCVE
Bos-taurus-UPF3A                        KLKKKDAKTGSIEDDPEYKKFLETYCVE
Canis-lupus-familiaris-UPF3A            KLKKKDAKTGSIEDDPEYKKFLETYCVE
Rattus-norvegicus-UPF3A                 KVKKKDAKTGSIEDDPEYKQFLESYSLE
Mus-musculus-UPF3A                      KLKKKDAKTGSIEDDPEYKQFLESYSLE
Tetraodon-nigroviridis-UPF3A            KLKKKDAKAGSIEEDPEYKRFLENISCD
Strongylocentrotus-purpuratus-UPF3B     IGKKVDARTATIEEDSDYKKFVETLNAE
Drosophila-melanogaster-UPF3            KARNDDSKVNTIESEPHYQEFIKRLAQE
Caenorhabditis-elegans                  NRMKEDTRVGAILTDKYYLDFCKKLEEE
Oryza-sativa-UPF3                       -NTKKDARQGTIMKDPEYLEFLESISKP
Arabidopsis-thaliana-UPF3               -SDKKDPREGSISKDPDYLEFLKVIAQP
Schistosoma-japonicum-UPF3A             KRDKVDKKQGSLLGDSEYIEFVKSMESA
                                           : * :  ::  :  *  * .
```

**Supplementary Figure S8. UPF3 alignments of NOPS-L region.** Homologous sequences obtained through submission of full-length human UPF3B isoform2 to NCBI BLAST (47) before aligning using EBI Clustal Omega multiple sequence alignment tool (48). The alignment is shown for human UPF3B residues 143-170, corresponding to the NOPS-L domain. An asterisk (*) indicates conserved residues in the alignment, a colon (:) indicates residues sharing conserved properties, while a period (.) indicates residues with weakly similar properties.

160 167

146 KRDTKVGTIDDDPEYRKFLESYATDNEKMTSTPETLLEEIEAKNRELIAKKTTPLLSFLK 205

206 NKQRMREEKREERRRREIERKRQREEERRKWKEEEKRKRKDIEKLKKIDRIPERDKLKDE 265

310

266 PKIKLLKKPEKGDEKELDKREKAKKLDKENLSDERASGQSCTLPKRSDSELKDEKPKRPE 325

326 DESGRDYREREREYERDQERILRERERLKRQEEERRRQKERYEKEKTFKRKEEEMKKEKD 385

396 399 403

386 TLRDKGKKAESTESIGSSEKTEKKEEVVKRDR 417

| Peptide | Identified PTM sites | Probability of PTM at each site |
|---|---|---|
| VGTIDDDPEYRK | Y10(Phospho) | T(3): 0.0; Y(10): 100.0 |
| FLESYATDNEK | Y5(Phospho) | S(4): 0.5; Y(5): 99.5; T(7): 0.0 |
| FLESYATDNEK | Y5(Phospho) | S(4): 6.3; Y(5): 93.2; T(7): 0.5 |
| LDKENLSDER | S7(Phospho) | S(7): 100.0 |
| LDKENLSDER | S7(Phospho) | S(7): 100.0 |
| LDKENLSDER | S7(Phospho) | S(7): 100.0 |
| LDKENLSDER | S7(Phospho) | S(7): 100.0 |
| LDKENLSDER | S7(Phospho) | S(7): 100.0 |
| ENLSDERASGQSCTLPK | S4(Phospho); | S(4): 100.0; S(9): 0.0; S(12): 0.0; T(14): 0.0 |
| ENLSDERASGQSCTLPK | S4(Phospho); | S(4): 100.0; S(9): 0.0; S(12): 0.0; T(14): 0.0 |
| ENLSDERASGQSCTLPK | S4(Phospho); | S(4): 100.0; S(9): 0.0; S(12): 0.0; T(14): 0.0 |
| KAESTESIGSSEK | S4(Phospho); S7(Phospho) | S(4): 89.3; T(5): 12.1; S(7): 98.6; S(10): 0.0; S(11): 0.0 |
| KAESTESIGSSEK | S4(Phospho); S7(Phospho) | S(4): 90.3; T(5): 9.8; S(7): 99.8; S(10): 0.0; S(11): 0.0 |
| KAESTESIGSSEK | S4(Phospho); S7(Phospho) | S(4): 91.1; T(5): 9.0; S(7): 99.9; S(10): 0.0; S(11): 0.0 |
| KAESTESIGSSEK | S4(Phospho); S7(Phospho) | S(4): 99.1; T(5): 1.0; S(7): 99.9; S(10): 0.0; S(11): 0.0 |
| KAESTESIGSSEK | S4(Phospho); S7(Phospho) | S(4): 89.3; T(5): 12.1; S(7): 98.6; S(10): 0.0; S(11): 0.0 |
| KAESTESIGSSEK | S4(Phospho); S7(Phospho) | S(4): 90.4; T(5): 19.1; S(7): 90.4; S(10): 0.0; S(11): 0.0 |
| KAESTESIGSSEK | S4(Phospho); S7(Phospho) | S(4): 98.9; T(5): 10.8; S(7): 90.4; S(10): 0.0; S(11): 0.0 |
| KAESTESIGSSEK | S4(Phospho); S7(Phospho) | S(4): 54.5; T(5): 54.5; S(7): 90.3; S(10): 0.5; S(11): 0.3 |
| KAESTESIGSSEK | S4(Phospho); S7(Phospho) | S(4): 50.4; T(5): 50.4; S(7): 99.1; S(10): 0.0; S(11): 0.0 |
| KAESTESIGSSEK | S4(Phospho); S7(Phospho) | S(4): 98.6; T(5): 12.1; S(7): 89.3; S(10): 0.0; S(11): 0.0 |
| KAESTESIGSSEK | S4(Phospho); S7(Phospho) | S(4): 98.6; T(5): 2.9; S(7): 98.6; S(10): 0.0; S(11): 0.0 |
| KAESTESIGSSEK | S4(Phospho); S7(Phospho) | S(4): 87.2; T(5): 25.5; S(7): 87.2; S(10): 0.0; S(11): 0.0 |
| KAESTESIGSSEK | S4(Phospho) | S(4): 90.3; T(5): 9.7; S(7): 0.0; S(10): 0.0; S(11): 0.0 |
| KAESTESIGSSEK | T5(Phospho); S7(Phospho) | S(4): 87.2; T(5): 25.5; S(7): 87.2; S(10): 0.0; S(11): 0.0 |
| AESTESIGSSEK | S3(Phospho); S6(Phospho) | S(3): 99.9; T(4): 97.8; S(6): 2.3; S(9): 0.0; S(10): 0.0 |
| AESTESIGSSEK | S3(Phospho); S6(Phospho) | S(3): 89.5; T(4): 20.9; S(6): 89.5; S(9): 0.0; S(10): 0.0 |
| AESTESIGSSEK | S3(Phospho) | S(3): 98.9; T(4): 1.1; S(6): 0.0; S(9): 0.0; S(10): 0.0 |
| AESTESIGSSEK | S3(Phospho) | S(3): 98.5; T(4): 1.5; S(6): 0.0; S(9): 0.0; S(10): 0.0 |
| AESTESIGSSEK | S9(Phospho) | S(3): 0.0; T(4): 0.0; S(6): 0.0; S(9): 9.8; S(10): 90.2 |
| AESTESIGSSEK | S3(Phospho); S6(Phospho) | S(3): 97.3; T(4): 5.3; S(6): 96.3; S(9): 0.5; S(10): 0.5; T(13): 0.0 |
| AESTESIGSSEK | S3(Phospho) | S(3): 98.0; T(4): 2.0; S(6): 0.0; S(9): 0.0; S(10): 0.0; T(13): 0.0 |

**Supplementary Figure S9. Phosphorylation mapping mass spectrometry of human UPF3B.** Data collected using purified full-size UPF3B expressed using the MultiBac insect cell expression system. The identified peptide fragments are highlighted in light blue, purple, orange, and red for peptides VGTIDDDPEYRK, FLESYATDNEK, ENLSDERASGQSCTLPK and AESTESIGSSEK, respectively. These peptides were identified as having multiples of +80 mass indicative of phosphorylation sites and a second fragmentation step identified residues with a more than 90% probability of harboring the phosphorylation.

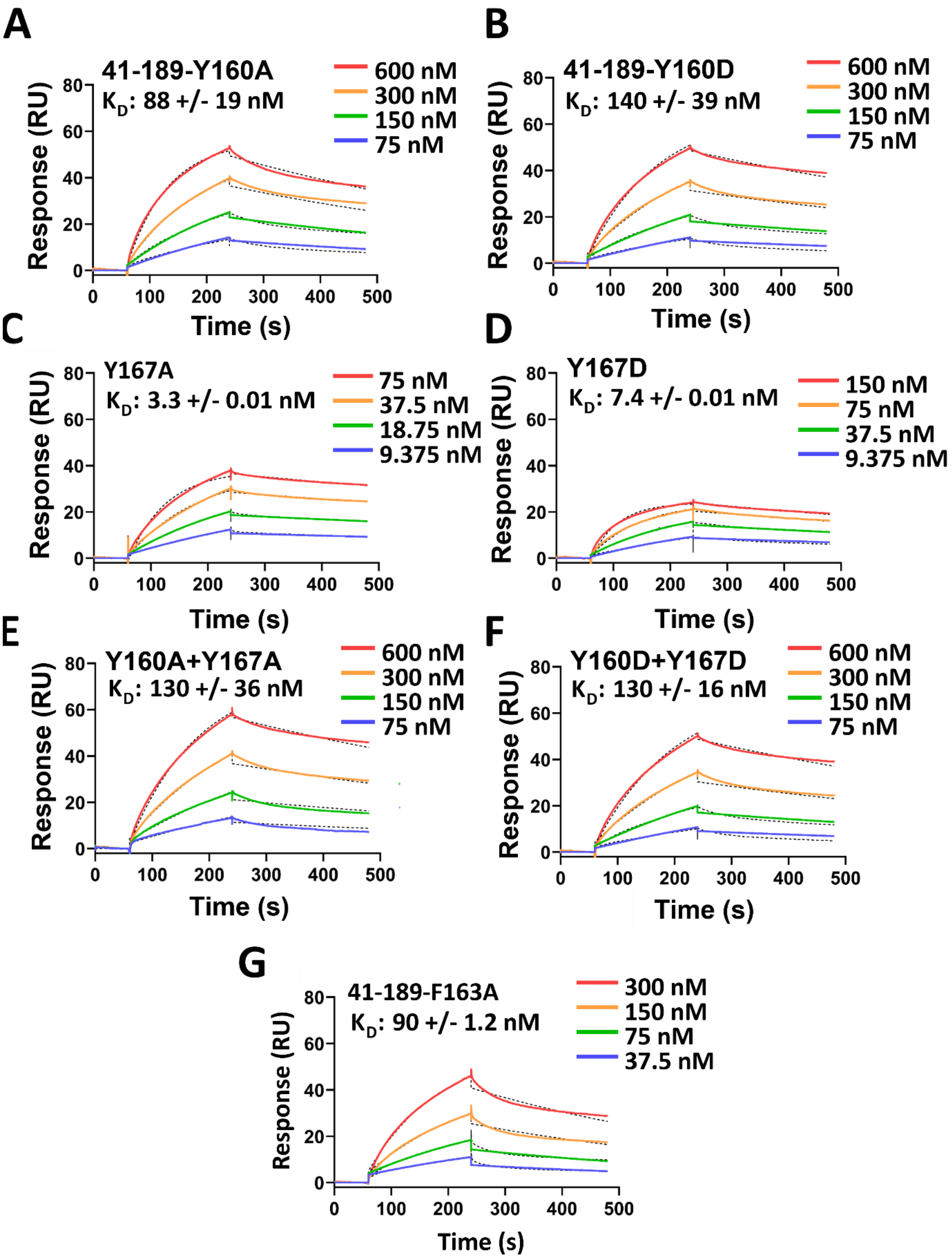
A
41-189-Y160A
$K_D$: 88 +/- 19 nM
600 nM
300 nM
150 nM
75 nM
Response (RU)
Time (s)
B
41-189-Y160D
$K_D$: 140 +/- 39 nM
600 nM
300 nM
150 nM
75 nM
Response (RU)
Time (s)
C
Y167A
$K_D$: 3.3 +/- 0.01 nM
75 nM
37.5 nM
18.75 nM
9.375 nM
Response (RU)
Time (s)
D
Y167D
$K_D$: 7.4 +/- 0.01 nM
150 nM
75 nM
37.5 nM
9.375 nM
Response (RU)
Time (s)
E
Y160A+Y167A
$K_D$: 130 +/- 36 nM
600 nM
300 nM
150 nM
75 nM
Response (RU)
Time (s)
F
Y160D+Y167D
$K_D$: 130 +/- 16 nM
600 nM
300 nM
150 nM
75 nM
Response (RU)
Time (s)
G
41-189-F163A
$K_D$: 90 +/- 1.2 nM
300 nM
150 nM
75 nM
37.5 nM
Response (RU)
Time (s)

**Supplementary Figure S10. SPR sensorgrams of UPF3B-41-189 variants binding to immobilized UPF2-MIF4GIII.** Representative sensorgrams for 4 analyte concentrations per UPF3B mutant are shown and their corresponding fits (black dotted lines). Fits were globally fitted with the 1:1 binding model within the T200 Biacore Evaluation Software to calculate indicated $K_D$ values. Sensorgrams are shown for UPF3B-41-189-Y160A **(A)**, UPF3B-41-189-Y160D **(B)**, UPF3B-41-189-Y167A **(C)**, UPF3B-41-189-Y167D **(D)**, UPF3B-41-189-Y160A+Y167A **(E)**, UPF3B-41-189-Y160D+Y167D **(F)**, and UPF3B-41-189-F163A **(G)**.

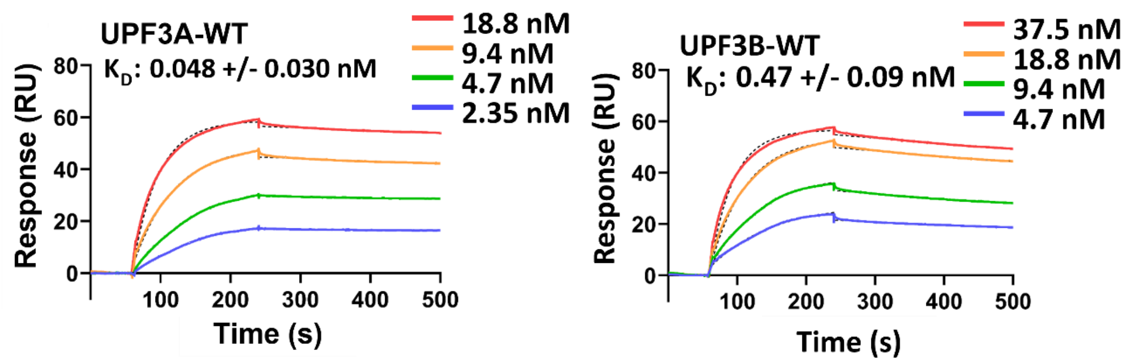


**Supplementary Figure S11. SPR sensorgrams of wildtype UPF3A and UPF3B analytes binding to immobilized UPF2-MIF4GIII.** Representative sensorgrams for 4 analyte concentrations of UPF3A-WT (left) and UPF3B-WT (right) binding to immobilized UPF2-MIF4GIII. Each concentration was individually fitted with the 1:1 binding model within the T200 Biacore Evaluation Software to calculate indicated $K_D$ values.

| Hex-Oligos | Sequence |
|---|---|
| HEX-24mer-ssDNA | 5’ HEX-CCC TGA GCT GAC GCA GCA CCT GGG 3’ |
| HEX-24mer-ssRNA | 5’ HEX-CAC UGA UCU GAC GCU GCA CCU GGG 3’ |
| HEX-24mer-dsDNA | 5’ HEX-CCC TGA GCT GAC GCA GCA CCT GGG<br>GGG ACT CGA CTG CGT CGT GGA CCC |
| HEX-24mer-dsRNA | 5’ HEX- CCC UGA GCU GAC GCA GCA CCU GGG<br>GGG ACU CGA CUG CGU CGU GGA CCC |

**Supplementary Table S1. HEX-oligo sequences used for fluorescence anisotropy and electrophoretic mobility shift assays.**

| Cloning Primers | Sequence |
|---|---|
| UPF3B.41.Forward | 5’ ACT CGC CAT GGA TCG CAA CAA GGA GAA GAA 3’ |
| UPF3B.143.Reverse | 5’ GGA CGT CGA CTT ACT TCT TTT TTG CAG CTT TTT G 3’ |
| UPF3B.262.Reverse | 5’ GGA CGT CGA CTT ATA ATT TGT CCC TTT CTG GA 3’ |
| UPF3B.189.Reverse | 5’ GGA CGT CGA CTT AAT TTT TTG CTT CTA TTT CCT CTA GC 3’ |
| UPF3B.214.Reverse | 5’ GGACGTCGACTTACTTTTCTTCTCTCATTCTCTGC 3’ |
| **Q5 Site Directed Mutagenesis Primers** | **Sequence** |
| UPF2-MIF4GIII-Avi-insert.Forward | 5' GCA GAA AAT TGA ATG GCA TGA AGA TTA CGA TAT CCC AAC G 3' |
| UPF2-MIF4GIII-Avi-insert.Reverse | 5' GCT TCA AAA ATA TCG TTC AGG CCG TGA TGG TGA TGG TGA TG 3' |
| UPF3B-Y160D.Forward | 5’ TGA TCC AGA AGA TAG AAA GTT TTT GG 3’ |
| UPF3B-Y160D.Reverse | 5’ TCA TCG ATA GTC CCG ACT TTG 3’ |
| UPF3B-Y160A.Forward | 5’ TGA TCC AGA AGC TAG AAA GTT TTT GGA AAG TTA TGC 3’ |
| UPF3B-Y160A.Reverse | 5’ TCA TCG ATA GTC CCG ACT TTG 3’ |
| UPF3B-F163A.Forward | 5’ ATA TAG AAA GGC ATT GGA AAG TTA TGC CAC 3’ |
| UPF3B-F163A.Reverse | 5’ TCT GGA TCA TCA TCG ATA G 3’ |
| UPF3B-Y167D.Forward | 5’ TTT GGA AAG TGA TGC CAC AGA CA 3’ |
| UPF3B-Y167D.Reverse | 5’ AAC TTT CTA TAT TCT GGA TCA TCA TC 3’ |
| UPF3B-Y167A.Forward | 5’ TTT GGA AAG TGC TGC CAC AGA CAA TG 3’ |
| UPF3B-Y167A.Reverse | 5’ AAC TTT CTA TAT TCT GGA TCA TC 3’ |
| UPF3B-Y160D-167D.Forward | 5’ TTG GAA AGT GAT GCC ACA GAC AAT GAG AAA ATG 3’ |
| UPF3B-Y160D-167D.Reverse | 5’ AAA CTT TCT ATC TTC TGG ATC ATC ATC GAT AG 3’ |
| UPF3B-Y160A-167A.Forward | 5’ TTG GAA AGT GCT GCC ACA GAC AAT GAG AAA ATG 3’ |
| UPF3B-Y160A-167A.Reverse | 5’ AAA CTT TCT AGC TTC TGG ATC ATC ATC GAT AG 3’ |

**Supplementary Table S2. Primers used for cloning of UPF3B truncation constructs (above) and UPF3B mutations and Avi-tagged variants (below).**
